# Benchmarking for genotyping and imputation using degraded DNA for forensic applications across diverse populations

**DOI:** 10.1101/2024.07.02.601808

**Authors:** Elena I. Zavala, Rori V. Rohlfs, Priya Moorjani

## Abstract

Advancements in sequencing and laboratory technologies have enabled forensic genetic analysis on increasingly low quality and degraded DNA samples. However, existing computational methods applied to genotyping and imputation for generating DNA profiles from degraded DNA have not been tested for forensic applications. Here we simulated sequencing data of varying qualities– coverage, fragment lengths, and deamination patterns–from forty individuals of diverse genetic ancestries. We used this dataset to test the performance of commonly used genotype and imputation methods (SAMtools, GATK, ATLAS, Beagle, and GLIMPSE) on five different SNP panels (MPS- plex, FORCE, two extended kinship panels, and the Human Origins array) that are used for forensic and population genetics applications. For genome mapping and variant calling with degraded DNA, we find use of parameters and methods (such as ATLAS) developed for ancient DNA analysis provides a marked improvement over conventional standards used for next generation sequencing analysis. We find that ATLAS outperforms GATK and SAMtools, achieving over 90% genotyping accuracy for the four largest SNP panels with coverages greater than 10X. For lower coverages, decreased concordance rates are correlated with increased rates of heterozygosity. Genotype refinement and imputation improve the accuracy at lower coverages by leveraging population reference data. For all five SNP panels, we find that using a population reference panel representative of worldwide populations (e.g., the 1000 Genomes Project) results in increased genotype accuracies across genetic ancestries, compared to ancestry-matched population reference panels. Importantly, we find that the low SNP density of commonly used forensics SNP panels can impact the reliability and performance of genotype refinement and imputation. This highlights a critical trade-off between enhancing privacy by using panels with fewer SNPs and maintaining the effectiveness of genomic tools. We provide benchmarks and recommendations for analyzing degraded DNA from diverse populations with widely used genomic methods in forensic casework.

**Highlights:**

- Biallelic SNP panels: >92% genotyping accuracy for 10X data with ATLAS
- Degraded DNA impacts accuracy under sequencing depth of 10X coverage
- Higher accuracies across genetic ancestries achieved with a diverse reference panel
- Leveraging population reference data is not applicable for small SNP panels
- Trade-off between genotype accuracy and privacy when considering SNP panel size

## 1. Introduction

The integration of next-generation sequencing (NGS) has opened new avenues of investigations within forensic genetics^1,2^. One area that has particularly benefited is human identification (HID) casework, which encompasses the identification of remains from mass disasters, military conflicts, cold cases, and historical identifications. HID cases often have relatively low success rates for generating DNA profiles due to the limited amount of DNA often recovered from these remains^3^. Over time DNA degrades, a process that may be accelerated due to environmental conditions, resulting in shorter DNA fragments, the accumulation of DNA damage such as deamination on the terminal ends of DNA strands, and an overall decrease in the amount of DNA present in the sample^4–6^. The complexities of degraded samples are further exacerbated by bacterial contamination that leads to lower endogenous content and in turn, lower depth of coverage. For this reason, it is often not possible to perform conventional short tandem repeat (STR) analysis, which requires longer fragments of DNA. However, NGS opens up the possibility of typing SNPs with lower quality samples. The challenges of working with degraded DNA in HID mirror those seen in ancient DNA analysis, where researchers analyze and extract DNA from archaeological specimens that are hundreds or thousands of years old^7^. Indeed, DNA profiles from degraded samples in forensic casework may show similar patterns of DNA degradation as ancient DNA, depending on environment, treatment, and storage conditions^8–10^. In the past decade, many new experimental and computational methods and tools have been developed to analyze degraded ancient specimens. The integration of ancient DNA laboratory methods has resulted in successfully generating DNA profiles from skeletal material (including burned remains)^10,11^ and hair^12^ in forensic casework. However, best practices for analyzing NGS data in general and degraded DNA data in particular, for forensic applications have not been developed.

Beyond the recovery and sequencing of genetic material from samples, the degraded nature of the recovered DNA also has implications for downstream analysis. This includes genome mapping, genotyping, imputation, and identification of individuals using genomic profiles. While there are clear analytical standards for DNA profiles following conventional forensic HID analyses (i.e. STR analysis from capillary electrophoresis data), there are no analogous standards for NGS of single nucleotide polymorphisms (SNPs) genotyping, as these analyses are currently typically used for non-courtroom applications such as generating investigative leads. Previous studies testing the impact of degraded DNA on different forensic NGS kits^13–15^ conflate the laboratory (i.e. library preparation and sequencing) and computational (mapping, genotyping, etc.) components of the workflow. A common observation of these studies is that incomplete DNA profiles lead to increased error rates for calling heterozygous sites. Probabilistic genotyping is commonly used in medical and population genetics and aims to incorporate sequencing error rates, allelic balance (i.e., proportion of reads supporting reference and non-reference alleles), and/or leverage population reference panels to improve genotype calling accuracy. In addition, a cost effective and scalable approach is to perform genotype imputation that leverages genetic information from population reference panels to infer genotypes from variants that were not directly typed or had low coverage^16^. However, the best practices and reliability of such methods remain unclear for use with degraded DNA from diverse populations. In particular, population reference panels are not equally representative for all human populations (as the vast majority of the individuals in genomic datasets have European-related ancestry^17^), potentially leading to unexpected biases across diverse worldwide groups.

In population and medical genetics studies, simulations and downsampling of high-coverage genomes has been used to test the accuracy of genotyping and imputation methods for low quality data^18–22^.

Typically, accuracy testing is performed on a subset of data or few ancestry groups from the dataset of interest to determine ideal methods and thresholds. However, given the real-world application of forensics analysis, it is essential to establish robust, validated workflows that can be widely applied. In addition, the impact of degradation on NGS analysis for diverse, worldwide populations has not been comprehensively tested for forensic applications.

As forensic investigations are focused on individual identification rather than population-based questions that directly impact present-day people around the world, understanding the limitations of genomic analyses is critical. Generalized guidelines for the analysis of NGS data including the impacts of DNA degradation on genotyping and imputation for individuals from diverse populations is necessary for forensics applications. This requires careful characterization of the false discovery rates for different parameters of each step of the sequence data analysis workflows, following laboratory testing. Here we present a simulation study where we examine how data quality (DNA fragment size, deamination patterns, and coverage) and variation in genetic diversity across diverse ancestry groups impacts various steps in genomic data analysis for five different SNP panels widely used in forensic or population genetics applications (**Figure 1**). We present benchmarks and limitations for each step, which may inform future forensic casework.

**Figure 1:**
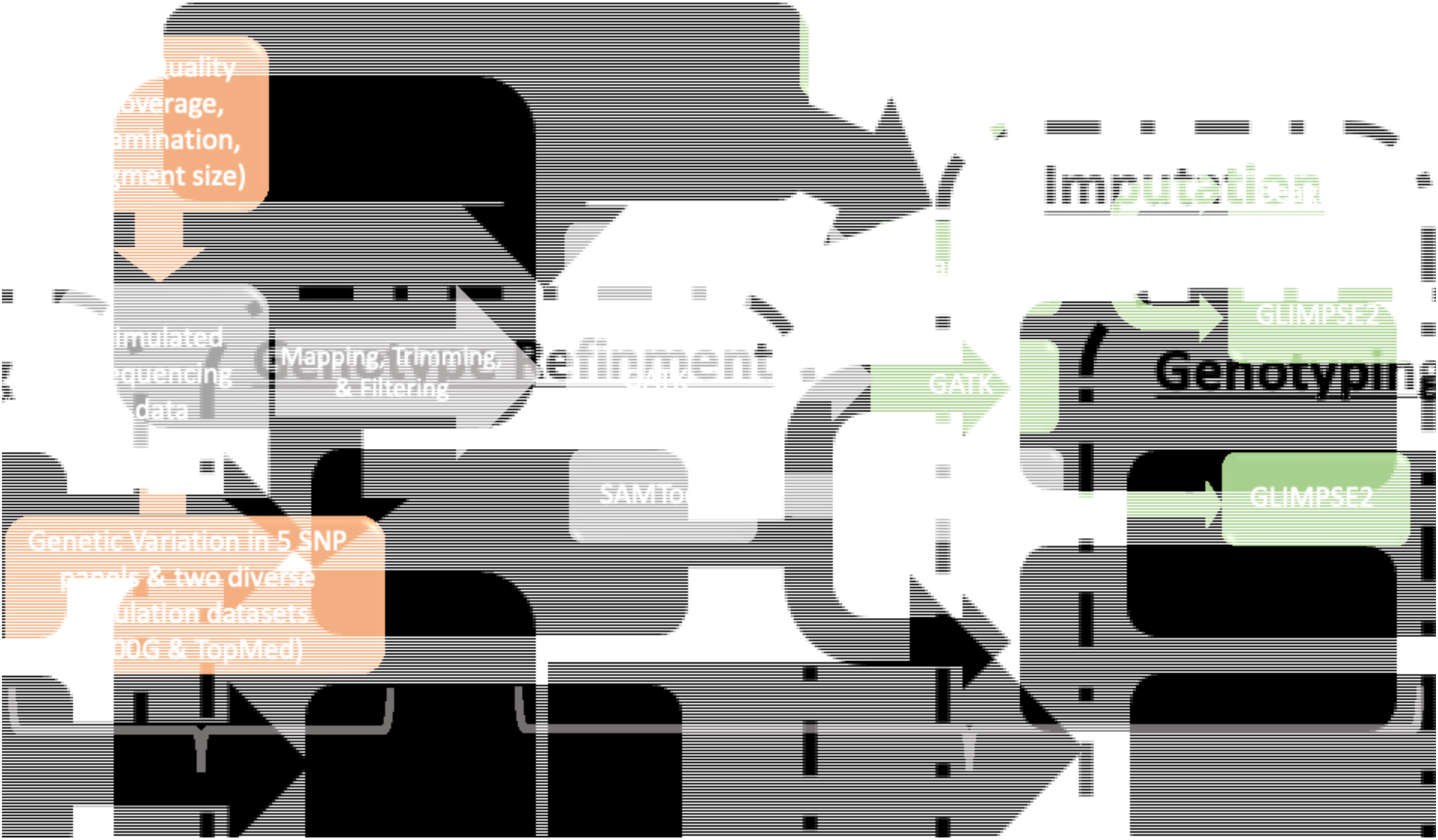
Overview of simulation workflow used in this study.

## 2. Materials and Methods

### 2.1 Simulated sequencing data

We used NGSNGS (version 0.9.2)^23^ to simulate single-end sequences of varying qualities from previously published DNA profiles^24,25^. These sequences were simulated using the human reference genome build hg38^26^, focusing on variants in four forensic and one population genetics SNP panels: FORCE^27^, MPS-plex^28^, the Gorden et al. 95K and 25K panels for extended kinship^29^, and the Affymetrix Human Origins array^30^. We used the publicly available genome sequences from 1000 genomes project^24^ and the TopMed dataset entitled NHLBI TopMed - NHGRI CCDG: The BioMe Biobank at Mount Sinai (phs001644.v3.p2) that was obtained through dbGAP (project #34943).

Genotypes were changed to reflect DNA profiles from ten individuals from the 1000 genomes project and thirty individuals from the TopMed study. The amount of overlap varied among SNP panels with the Gorden 95K and 25K having the most shared SNPs with other panels (48% and 65% respectively), the Human Origins array and MPS-plex with the least shared SNPs (6.7% and 4.5% respectively) and the FORCE panel in the middle (23%) (**Figure 2A**). We removed insertion-deletions (indels) that overlapped any SNP positions, resulting in a range of 1,209 to 600,706 autosomal SNPs per panel (**Table S1**).

**Figure 2.**
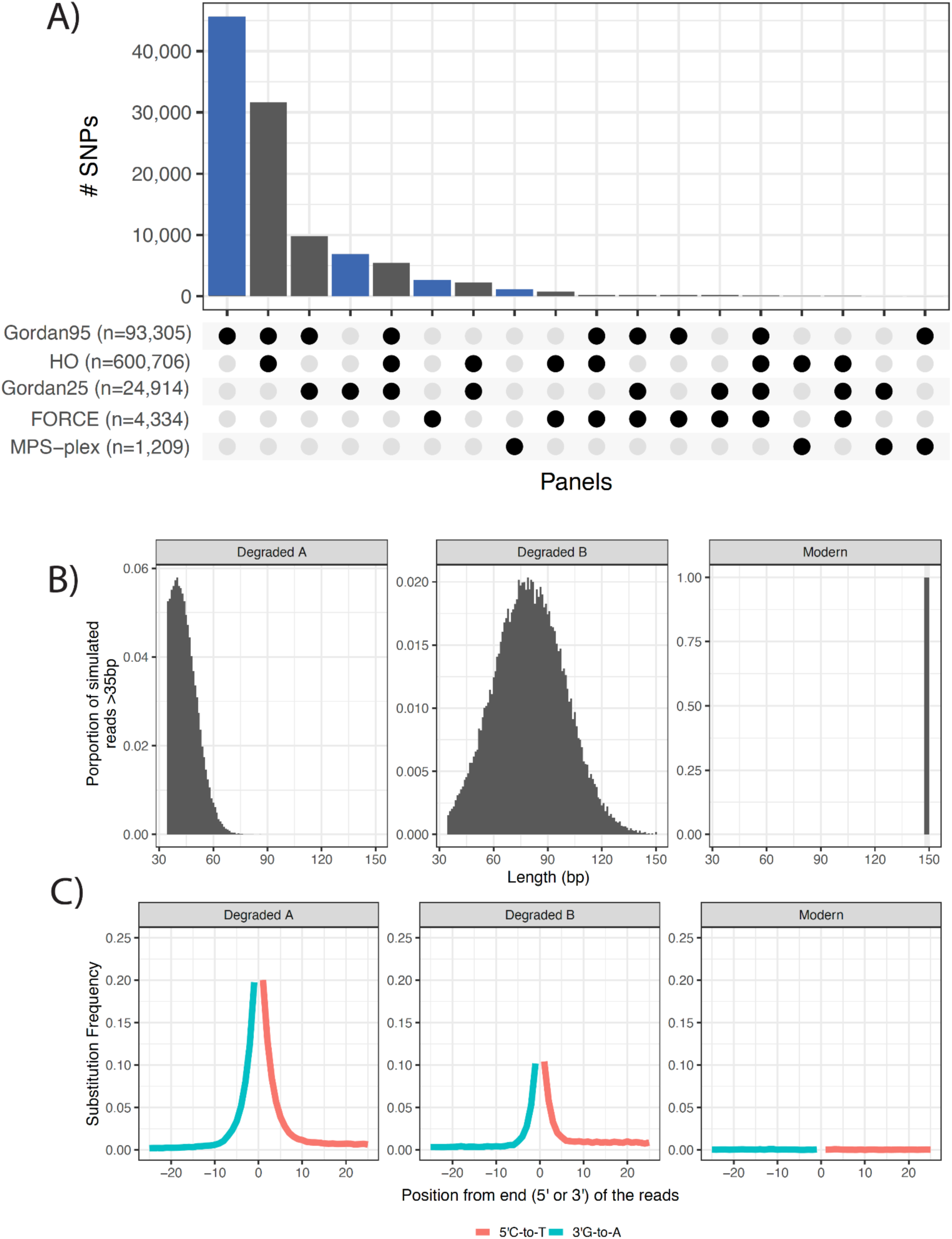
Parameters used for simulating sequencing data. A) Number of unique or overlapping SNPs between panels after removing indels used for introducing genetic variation from published genetic data. Colored bars represent SNPs unique to a respective SNP panel. Note: 560,175 SNPs unique to the Human Origins panel are not shown. B) Fragment length distributions and C) putative deamination substitution patterns used for simulating three variations of sequence data.

We simulated three sets of sequences, one matching the characteristics of modern DNA and two with varying amounts of degradation. For modern simulations, sequences of 150 base pairs (bp) were generated with no deamination. For the degraded simulation variation A, sequence lengths follow a normal distribution with an average length of 40bp and variance of 10 (NGSNGS option: -ld norm,40,10), with deamination added following the Brigg’s model (Briggs et al., 2007) (NGSNGS option: -m b7,0.024,0.36,0.68,0.0097) (**Figure 2B**). For degraded simulation variation B, we simulated sequences that are less degraded with sequenced read lengths that follow a normal distribution around 80bp with variance of 20 (NGSNGS option: -ld norm,80,20) with reduced deamination (NGSNGS option: -m b7,0.01,0.52,0.42,0.0097) compared to variation A (**Figure 2C**). For reproducibility, a seed value of 22688 was used for simulations. Each set of sequences was simulated to coverages of 0.1X, 0.5X, 1X, 5X and 10X across the genome. Quality scores and sequencing errors were generated using a matrix for the Illumina HiSeq 2500 platform from the NGSNGS package (AccFreqL150R1.txt).

### 2.2 Genotype data, population definitions, and population reference panels

In order to test the impact of genetic ancestry on the accuracy of genotyping and imputation methods, we simulated sequencing data from individuals from the 1000 genomes project^24^ and the TopMed study. We included the latter as the individuals are more reflective of the diversity within the United States. We used genotypes from ten individuals in the 1000 genomes dataset for simulations, including two individuals from each of the five superpopulations defined for the 1000 genomes dataset (European, African, American, South Asian, and East Asian)^31^. We used genotypes from thirty individuals within the TopMed study including individuals that self-identified as African-American/African, East/South-East Asian, European American, Hispanic/Latin American, South Asian, or multiple/other designations. Below we describe our selection criteria.

In human genetic studies, it is common to present results by groups, based on ancestry or geographic location, etc. However, these groups may not reflect the continuous nature of the genetic ancestries of individuals, in particular for people of mixed genetic ancestry^32^. Such grouping can also present a challenge when testing accuracies for genotype refinement and imputation, as these methods rely on population reference panels. In this case reference group labels may not capture the relationship between the individual of interest and the selected population reference panel, in turn leading to biases and reduced accuracy of genotype calls. To assess the impact of choice of reference population, we explored how use of different population reference panels composed of individuals of different ancestry or continental groups can impact genotype refinement and accuracy, as a function of genetic similarity to the individual of interest. We used the widely used statistic in population genetics, *FST*, to measure the genetic similarity of each individual to each of the geographical superpopulations used as population reference panels. We note estimates of *FST* using a single diploid genome can be very noisy and sensitive to ascertainment bias (choice of SNPs used for the analysis) and hence our estimates should be considered as a rough estimate of genetic similarity and not as an evaluation of population structure. *FST* was calculated by performing *smartpca*^33^ with a simulated individual from either ten individuals from 1000 genomes (only a subset of 1000 genomes was used as this dataset was used for creating population reference panels) or 11,996 individuals from TopMed study. Thirty individuals from the TopMed study were then selected to cover the distributions of observed *FST* values in relation to the different 1000 Genomes superpopulations (African, American, East Asian, South Asian, and European) and a compilation of individuals from all five superpopulations (**Figure 3**).

**Figure 3:**
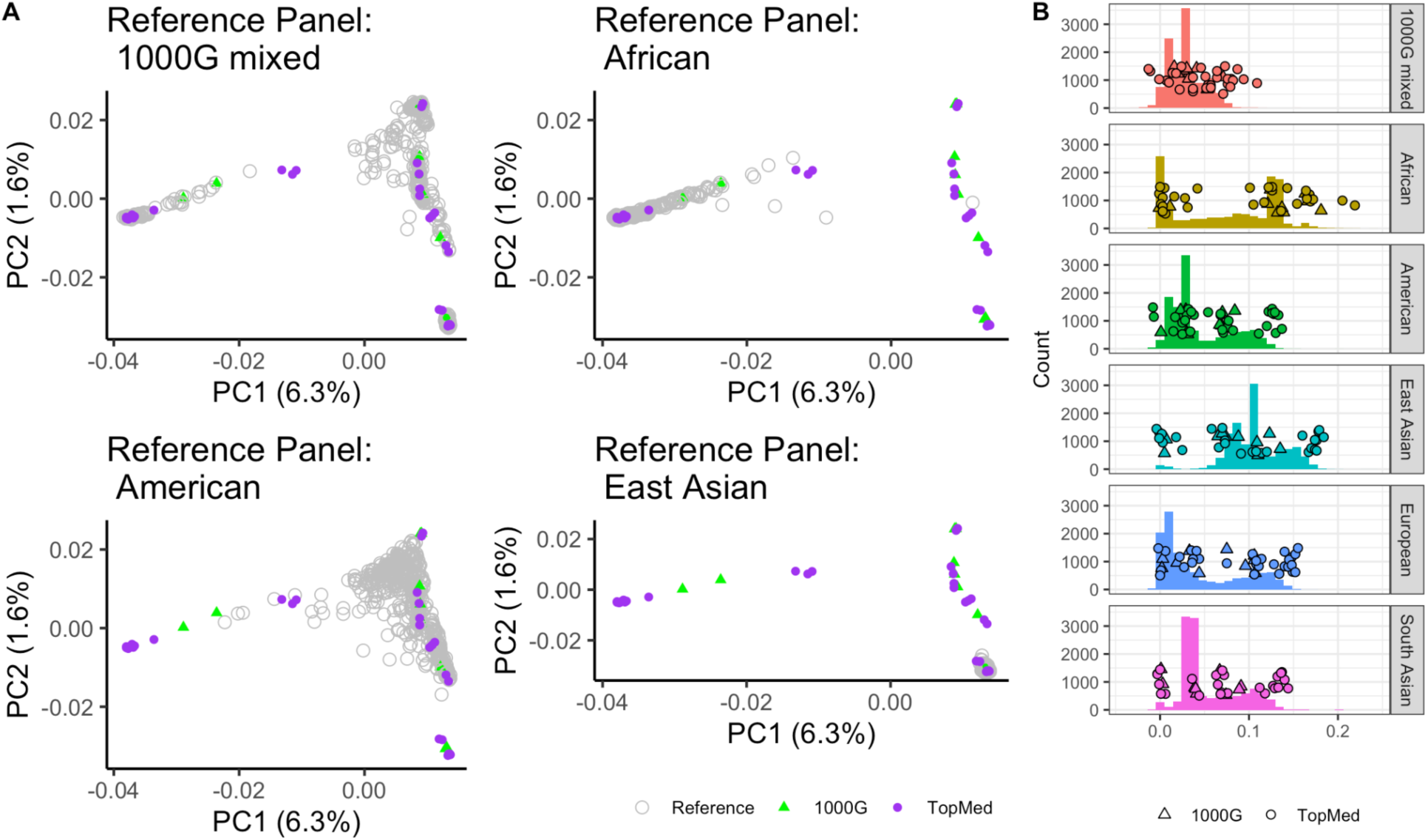
Genetic similarity between individuals used for simulations and selected population reference panels. A) PCA of four selected population reference panels, each with 450 individuals from the 1000 genomes dataset (shown in gray), and the forty individuals (shown in green (1000 Genomes) and purple (TopMed)) used for simulations. PCA was generated using smartpca^30^. B) Histogram of calculated *FST* values between individuals in the TopMed study and six potential population reference panels. Points on top of the histogram represent the forty (thirty TopMed individuals and ten 1000 genomes) individuals selected for simulations in this study.

Based on the results of the *FST* tests, we selected four population reference panels for testing in this study. Two (1000 Genomes African and East Asian) showed the greatest genetic dissimilarity in relation to the individuals tested (**Figure 3**). One (1000 Genomes American) represents a more admixed population and one (1000G mixed) represents more genetic diversity as it includes individuals from all super populations in this larger dataset and resulted in the narrowest range of *FST* values (**Figure 3**). European and South Asian were not included to reduce the number of parameters tested and because the *FST* values observed with these population reference panels were within the ranges observed for the African, East Asian, and American superpopulations. The three super population panels each comprise 450 individuals that were randomly selected from each respective super population, excluding individuals used within the simulation. The 1000G mixed population reference panel was constructed by selecting 90 random individuals from each of the five super populations (450 in total), also excluding individuals used for the simulation.

### 2.3 Processing of sequencing data

The simulated sequencing reads were mapped to the human reference genome build (hg38) using bwa aln ^34^ with the modified parameters typically used for ancient DNA sequencing data^35^. This includes disabling the seed for the alignment (option -l 16500), increasing tolerance for differences from the reference genome (option -n 0.01) and increasing the maximum number of gaps (option -o 2). After mapping, we used SAMtools (version 1.8) ^36,37^ to filter the reads, retaining only reads of at least 35 bp in length with a mapping quality of at least 25. Finally, to limit the impact of deamination, C-to-T and G-to-A substitutions on the terminal 8 bp of the reads were masked (**Figure S3**).

### 2.4 Concordance Evaluations

Concordance was evaluated by calculating the number of correctly genotyped SNPs over the total number of genotyped SNPs for each set of parameters (coverage, DNA quality, SNP panel, and chromosome). For heterozygous sites, we did not phase the genotypes and thus considered 0/1 or 1/0 as concordant. The following equation was used to calculate heterozygosity per individual for each panel and chromosome combination:

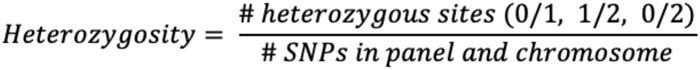

In order to compare different methods directly, we also calculated relative concordance rates between methods (i.e. genotype methods, genotype refinement, or imputation). For each individual, chromosome, DNA quality, coverage, and SNP panel variation the method with the highest number of concordant SNPs was identified (max concordant SNPs). We considered the ratio of the number of concordant SNPs for each method and the max concordant SNPs resulting in the relative concordance rate which was compared across individuals (**Table 1** for example).

**Table 1.**
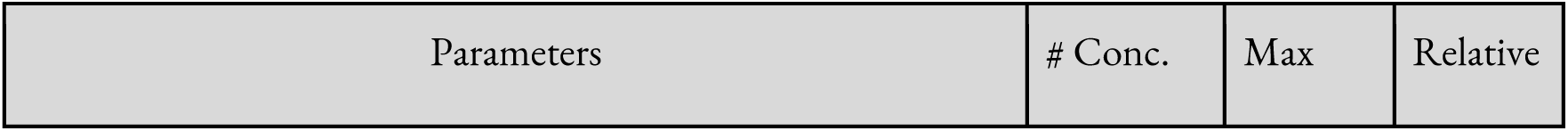
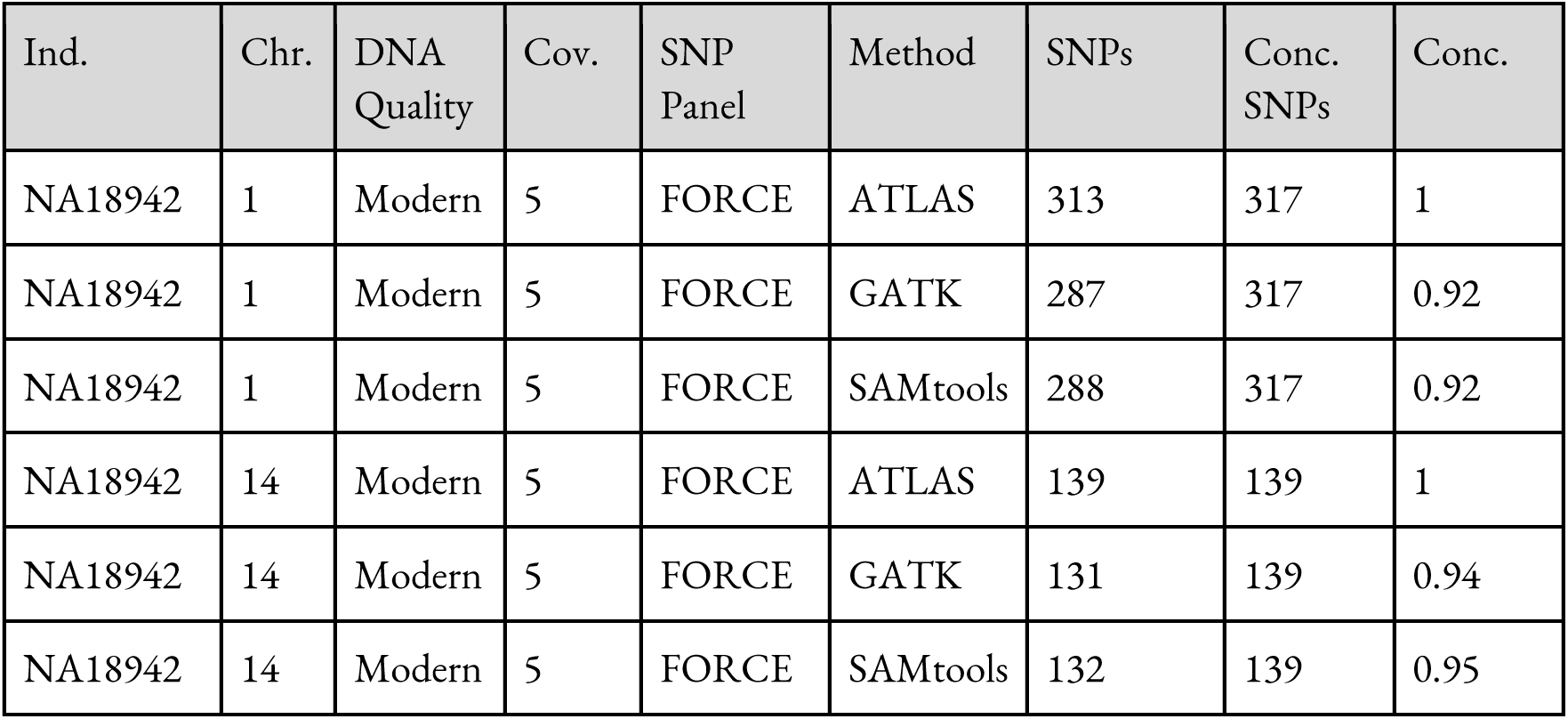
Example of relative concordance calculation. Each column shows a step in the calculation. Ind. = Individual; Chr. = Chromosome; Cov. = Coverage; Conc. = Concordant/Concordance

| Ind. | Chr. | DNA Quality | Cov. | SNP Panel | Method | SNPs | Conc. SNPs | Conc. |
| --- | --- | --- | --- | --- | --- | --- | --- | --- |
| NA18942 | 1 | Modern | 5 | FORCE | ATLAS | 313 | 317 | 1 |
| NA18942 | 1 | Modern | 5 | FORCE | GATK | 287 | 317 | 0.92 |
| NA18942 | 1 | Modern | 5 | FORCE | SAMtools | 288 | 317 | 0.92 |
| NA18942 | 14 | Modern | 5 | FORCE | ATLAS | 139 | 139 | 1 |
| NA18942 | 14 | Modern | 5 | FORCE | GATK | 131 | 139 | 0.94 |
| NA18942 | 14 | Modern | 5 | FORCE | SAMtools | 132 | 139 | 0.95 |

### 2.5 Genotyping and genotype refinement

Three different genotyping methods, paired with genotype refinement, were tested in this study that have previously been used for forensic or ancient DNA genotyping. For each step that required a reference genome the human reference genome build (hg38) was used. The resulting vcf files were then intersected with the SNP panels of interest using BEDtools (version 2.28.0)^38^ on the relevant chromosome (1 or 14).

1) ATLAS (version 0.9)^21^ was used with the maximum likelihood genotype caller option as recommended in Hui et al., 2020^18^. Beagle 4.1^39^ was used for genotype refinement with default parameters for each of the four selected population reference panels from the 1000 genomes dataset. For this analysis, the impute option was set to false and gprobs to true. Positions with a resulting genotype probability of at least 0.99 were retained.
2) GATK (version 4.1.9.0)^40^ HaplotypeCaller following the best practices recommendations^41,42^. For genotype refinement GATK’s CalculateGenotypePosteriors was used with default settings for each of the four selected population reference panels (option -supporting). BCFtools (version 1.6)^36^ was then used to retain only positions with a genotype quality (GQ) of at least 20.
3) SAMTools (version 1.8)^36,37^ mpileup with the standard recommendations with an adjusted mapping quality of 50 (option -C50), readjusting base quality scores with a minimum of 25 (options --redo-BAQ --min-BQ 25) with genotype likelihoods generated in BCF format (option --BCF). The output was then piped directly into BCFtools (version 1.6) for consensus calling (option --consensus-caller). SAMTools is not typically paired with population reference

panels for genotype refinement so we performed refinement by using BCFtools to only retain positions with a quality (QUAL) of at least 20.

### 2.6 Imputation

We tested two different imputation methods– Beagle5.4 and GLIMPSE2– using three panels of SNPs (25K, 95K, and HO) with simulated data of at least 0.5X coverage. For each analysis, the VCF generated using ATLAS + Beagle 4.1 workflow described above (Methods 2.4 and 2.5) was used. We used the four reference panels from the 1000 Genomes described above for genotype refinement. In addition, the VCF generated from SAMtools post-genotype refinement (Methods 2.4 and 2.5) was also used with GLIMPSE2.

1) Beagle 5.4^43^ was used with default parameters (except gp=true) using the same population reference panel as used for genotype refinement. Only positions with a genotype probability of at least 0.99 were retained and BEDtools was then used to subset the output to only include the SNPs on the SNP panel of interest.
2) GLIMPSE2 ^44,45^ (v2.0.0). The estimated genotype probabilities from the output of Beagle 4.1 were converted to Phred scaled genotype likelihoods with BCFtools^37^ +tag2tag. This VCF was then input into GLIMPSE2 ^44, 45^ for phasing and imputation using each of the same population reference panels as used above. For creating the binary reference chunks the sequential model was used with 4Mb chunking and the GLIMPSE2 provided genetic map for hg38 as recommended. The resulting output was then filtered to retain only positions with a genotype probability of at least 99%.

## 3. Results

### 3.1 Simulation framework

We used NGSNGS^23^ to simulate single end sequencing data from forty individuals with varying data qualities– modern, degraded A, and degraded B (see **Methods**, **Figure 2B-C**). The two sets of degraded DNA differ in data quality due to variability in deamination rates and fragmentation patterns. Degraded A reflects DNA damage patterns similar to what has been observed in highly degraded forensic and ancient DNA samples^8–10,46^. Degraded B has lower levels of deamination and fragmentation (**Figure 2**), which is presumed to be observed more often in degraded forensic samples.

In turn, modern reflects high quality and complete DNA sequences with no deamination to provide a benchmark for comparisons. These data were used to examine the impacts of genome mapping and genotyping for five SNP panels commonly used in forensics and ancient DNA studies (**Figure 2A; Supplementary Section 1**). We used SNP panels, rather than low-coverage sequencing data, as they are more commonly used in the forensic field due to greater privacy concerns, efforts to maintain consistency with previously validated workflows, and to reduce costs^47,48^. The five SNP panels tested include: (a) the MPS-plex panel (1,270 SNPs ^28^) that consists of tri-allelic loci and was developed to increase power for individualization with less SNPs for application with degraded DNA, (b) the FORCE panel (5,422 SNPs^27^) that was developed as an “all-in-one” panel for wide-spread use in the forensic field and covers loci informative for phenotyping, genetic ancestry, kinship and individualization, (c-d) the 95K and 25K panels^29^ were developed for identifying distant kinship (up to 4th degree relatives with 94,752 and 24,999 SNPs respectively) and (e) the Affymetrix Human Origins array^30^ that contains 1.2 million SNPs ascertained in diverse populations and is widely used in ancient DNA and population genetics studies. We generated sequencing data from the profiles of forty individuals, ten from the 1000 genomes high coverage dataset^24^ and thirty from TopMed^25^ study, to cover a range of genetic ancestries (see **Methods** for selection of individuals).

### 3.2 Sequence mapping and trimming for degraded DNA

The first step in genome sequence analysis is to map reads to the human reference genome. We used the simulated sequences (as described above) and mapped them to the human reference genome build hg38^26^ using one of the most widely used methods, *bwa*^34^. We considered two options– the standard options for modern DNA and another with more relaxed parameters proposed for ancient DNA. For ancient genomes, it has been previously shown that disabling the seed region and increasing permissibility around the number of allowed gaps and differences from the reference genome increases the number of mapped degraded sequences^35^. We found that the relative ratio of reads between the two settings was similar (<1.006) (**Supplementary Section 2**). However, the small differences translate to retention of thousands of reads using the ancient DNA parameters even for coverages of 0.1X (**Figure S1**). This suggests that using ancient DNA mapping parameters can be helpful to retain more mapped DNA sequences and hence we use this pipeline for the rest of the analyses described below.

Another step in genome mapping is to remove adapter sequences and trim the terminal bases of the reads to remove either low quality bases (modern DNA analyses^49,50^) or deamination (ancient DNA^51^). For modern data trimming is typically performed before mapping (at fastq level), while for ancient DNA data trimming for deamination is performed after mapping (at bam level). The difference in setups is motivated by the observation that the presence of deamination can be used for ancient DNA authentication^5^. However, trimming short reads imposes a trade-off between loss of data (especially as reads are only 80-100 bp long in ancient DNA) and mitigating impact of DNA damage by removing low quality bases. Thus, we evaluated the impact of trimming terminal ends at either the pre-mapping or post-mapping (**Figure S3**) by comparing the genotyping accuracy for transitions in degraded DNA (**Supplementary Section 1.3**). We find that trimming pre-mapping (at the fastq level) results in significantly less concordance for low coverage sequences (1X to 5X; **Table S3, Figure S4**).

Surprisingly, trimming after mapping does not have a significant advantage compared to not trimming at all until higher coverages (>5X; **Table S3, Figure S4**). As masking of putative deamination on the terminal ends of reads post-mapping (at the bam level) is not detrimental, we performed this step for all subsequent analyses.

### 3.3 Genotyping Accuracy

We evaluated the impact of DNA quality on the accuracy of genotyping using three different genotypers commonly used for ancient DNA and forensic applications: ATLAS^21^, GATK^40–42^, and SAMtools^36,37^. To this end, we simulated sequencing data for forty individuals from diverse populations for two chromosomes, 1 and 14 (the largest chromosome and a mid-size chromosome) (see **Methods**). The resulting sequences were mapped to the human reference genome (Hg38) using *bwa* with ancient DNA parameters (see **Methods**). We filtered the mapped reads based on length and mapping quality (≥35 bp long and with mapping quality of ≥25) with masking for deamination on the terminal ends of reads and used the resulting sequences as the input to each of the genotypers. The output VCF was then subset to variants on each of the five SNP panels (see **Methods**). We calculated the concordance as the number of genotyped SNPs that match the original data out of all the SNPs genotyped on each SNP panel (we note, for heterozygous sites, we do not consider phase so 0/1 and 1/0 are considered concordant genotypes) (**Figure 4**). As expected, we find higher coverage results in greater concordance across all SNP panels. The smallest SNP panels (FORCE and MPS-plex) have greater variance in concordance than other panels likely due to the relatively smaller number of SNPs on chromosomes 1 and 14 in these two panels (150-347 and 35-100 SNPs respectively).

**Figure 4:**
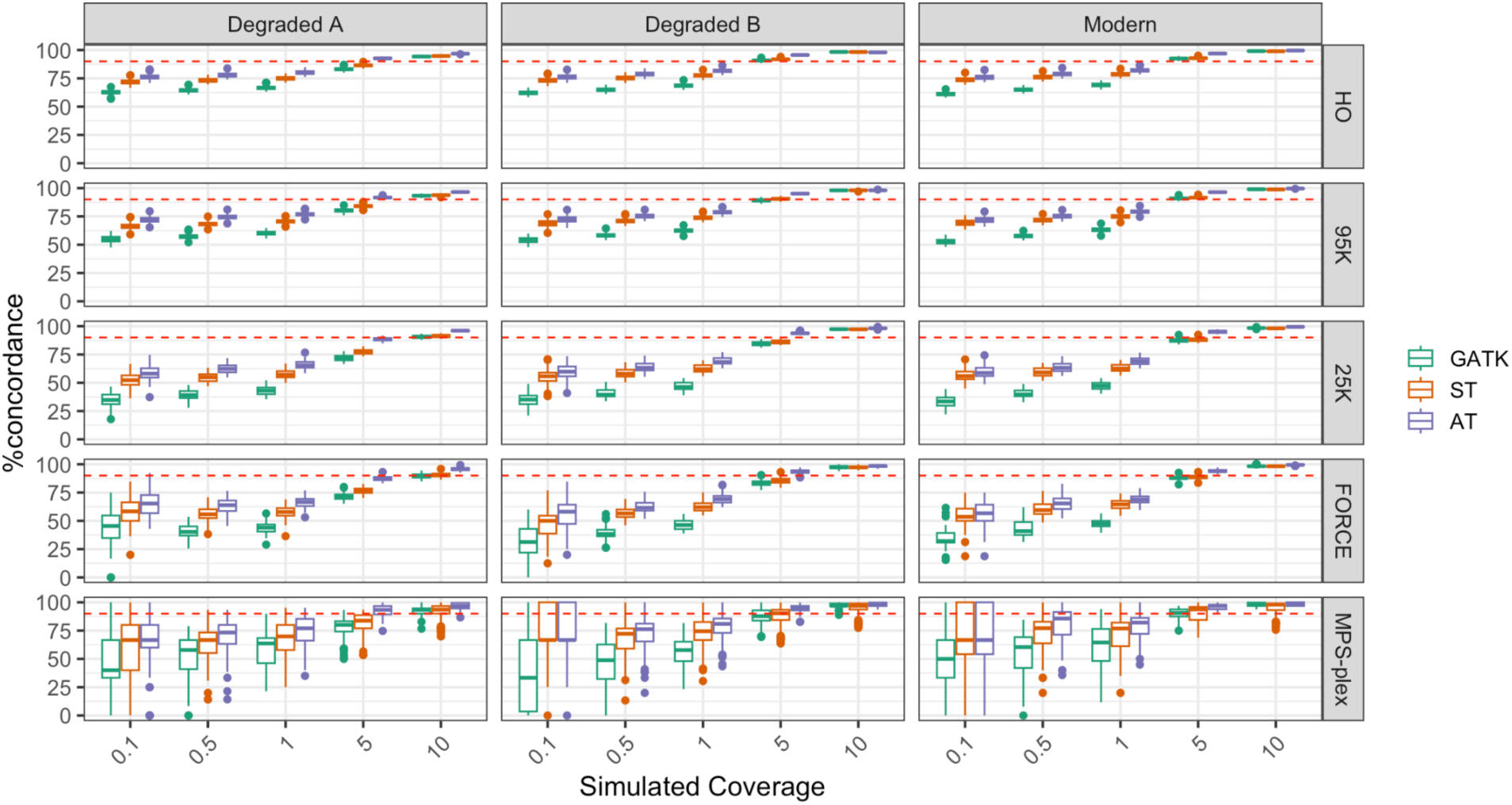
Concordance of genotypes for different genotyping methods. Each coverage, DNA quality, genotyper, and SNP panel combination is represented by simulated sequencing data from 40 individuals. The Tukey box and whisker plots show the lower and upper quartiles boxed with the whiskers representing 1.5 times the interquartile range. Dots denote outliers. The red dashed line denotes 90% concordance.

Heterozygous sites tend to have higher error rates due to allelic dropout, misalignment or base calling errors that are exacerbated in low quality or low coverage data. Therefore, we evaluated the concordance with the ATLAS genotyper at sites where the expected genotype was heterozygous (i.e. 0/1, 0/2, or 1/2 considering triallelic loci), for resulting DNA profiles with more than 5 heterozygous sites. We find that at low coverages concordance for heterozygous sites is low, such that reaching a concordance rate of >50% requires at least 5X coverage depth for modern DNA and degraded B data qualities (**Figure S5**), and 10X coverage depth for degraded A DNA quality. Notably, if only 0/1 heterozygous sites (i.e., excluding triallelic sites) are examined across the four largest SNP panels (excluding the smallest panel MPS-plex), minimum concordance rates increase from 55.6% and 80% to 62.5% and 90.2% for 5X and 10X respectively (**Figure S6**). This is compared to concordance rates of >92% for homozygous sites at 5X coverage of the four largest SNP panels. As there are significant differences in the genome-wide heterozygosity (i.e., proportion of heterozygous sites across the genome) among human populations^31^, we assessed the impact of an individual’s heterozygosity on the observed concordance rates across sequence coverages. For the SNPs on each SNP panel, we estimated the rate of heterozygosity (i.e. proportion of sites that were heterozygous for each individual in relation to the number of ascertained SNPs on each SNP panel per chromosome) and compared this estimate to the observed concordance in genotypes (**Figure S7**). We find that increased heterozygosity correlates with decreased concordance for coverages less than 10X (Pearson correlation, max p-value 0.047 and min correlation -0.23, **Figure S7**). Moreover, lower minor allele frequencies (MAF) have previously been shown to coincide with decreased non-reference concordance^22^. Therefore, we also examined this impact. Given that we examined SNPs on genotyping arrays, we found relatively few variants with MAFs <5% (**Figure S8**). The overall concordance (including 0/0, 0/1 and 1/1 genotypes) is negatively correlated to MAF, particularly at low coverages and proportional to the fraction of homozygous non-reference alleles (**Figure S8 and S9**). Heterozygous concordance was not observed to correlate with MAF (**Figure S10**), but this may be due to a lack of power given the low number of rare variants.

Next, we compared the concordance across the three genotypers, ATLAS, SAMtools and GATK, by calculating the relative ratio of concordance estimated for each method (using the best practices and parameters recommended by each method, see **Methods**). The relative concordance was calculated by identifying the genotype method with the highest number of concordant SNPs for each setting, where we vary the DNA quality, SNP panel, coverage, and chromosome used. Thus, the best setting for a set of parameters yields a relative concordance rate of 1. ATLAS resulted in consistently higher concordance across coverages, DNA qualities, and SNP panels (**Figure S11**). Simulated modern, 10X coverage data resulted in the lowest error rates across genotypers (median error rates across SNP panels 0.4-2%, 0.9-2%, and 1.1-2% for ATLAS, GATK, and SAMTools respectively) and simulated degraded A, 0.1X coverage data had the highest error rates (median error rates across SNP panels 24-33%, 37- 65%, and 28-48%for ATLAS, GATK, and SAMTools respectively). As ATLAS was developed specifically for genotyping low-quality, degraded sequencing data, this result is not surprising, but emphasizes its applicability for forensic analysis.

### 3.4 Genotype Refinement

After genotyping, it is typical to infer genotype probability or likelihoods to enhance the quality of genotype calls and flag potential errors. For instance, one approach is to use population reference panels to determine the likelihood of observing a particular genotype in the reference dataset. Another approach, recommended by SAMtools, is to refine the outputs based on the computed quality scores (**Methods 2.4**) Following each method’s recommendations, we performed genotype refinement for each genotyper. For ATLAS and GATK, we performed genotype refinement using four population reference panels of equal size (450 individuals) that were subsets of individuals from the 1000 genomes project high coverage data, phase 4^24^. Three of these population reference panels were from the five superpopulations defined in the 1000 genomes project (African, American, and East Asian, see **Methods**) and the fourth consisted of an equal mixture of individuals from all five superpopulations (90 individuals from each super population) (**Figure 3**). For GATK, we followed the variant recalibration best practices and we used Beagle 4.1^39^ for ATLAS (**Methods 2.4**).

We find genotype refinement results in a significant change in genotype concordance for almost all SNP panel and coverage combinations across genotypers (max p-value for coverages 1-10X: 6.8E-4, Wilcoxon test with Bonferroni correction for multiple testing; **Table S4**). For the three largest SNP panels this is due to an increase in accuracy with genotype refinement, however we did observe reduced accuracy in some instances for coverages ≤0.5 for the two smallest SNP panels (**Figure S12**). More specifically, post refinement with ATLAS + Beagle 4.1 results in a median concordance rate of >88% at 0.1X for the three largest SNP panels (Human Origins, 95k, and 25K) (**Figure S13**). For comparison, similar levels of genotype concordance were only obtained with coverages of at least 5X without genotype refinement (**Figure 4**). As expected, refinement increases specificity at the cost of sensitivity. Post-refinement we observed a substantial decrease in the total number of SNPs. For example, at 5X coverage for the 25K panel with ATLAS genotyping followed by refinement with Beagle 4.1, we observed a decrease of 58%, 49%, and 48% in the proportion of SNPs covered on average for degraded A, degraded B, and modern DNA qualities (**Figures S13-14, Figure 5**). Notably, ascertainment panels with more SNPs resulted in higher accuracies, likely as this allows the methods to leverage allelic correlation patterns across neighboring SNPs in the population reference panels.

**Figure 5.**
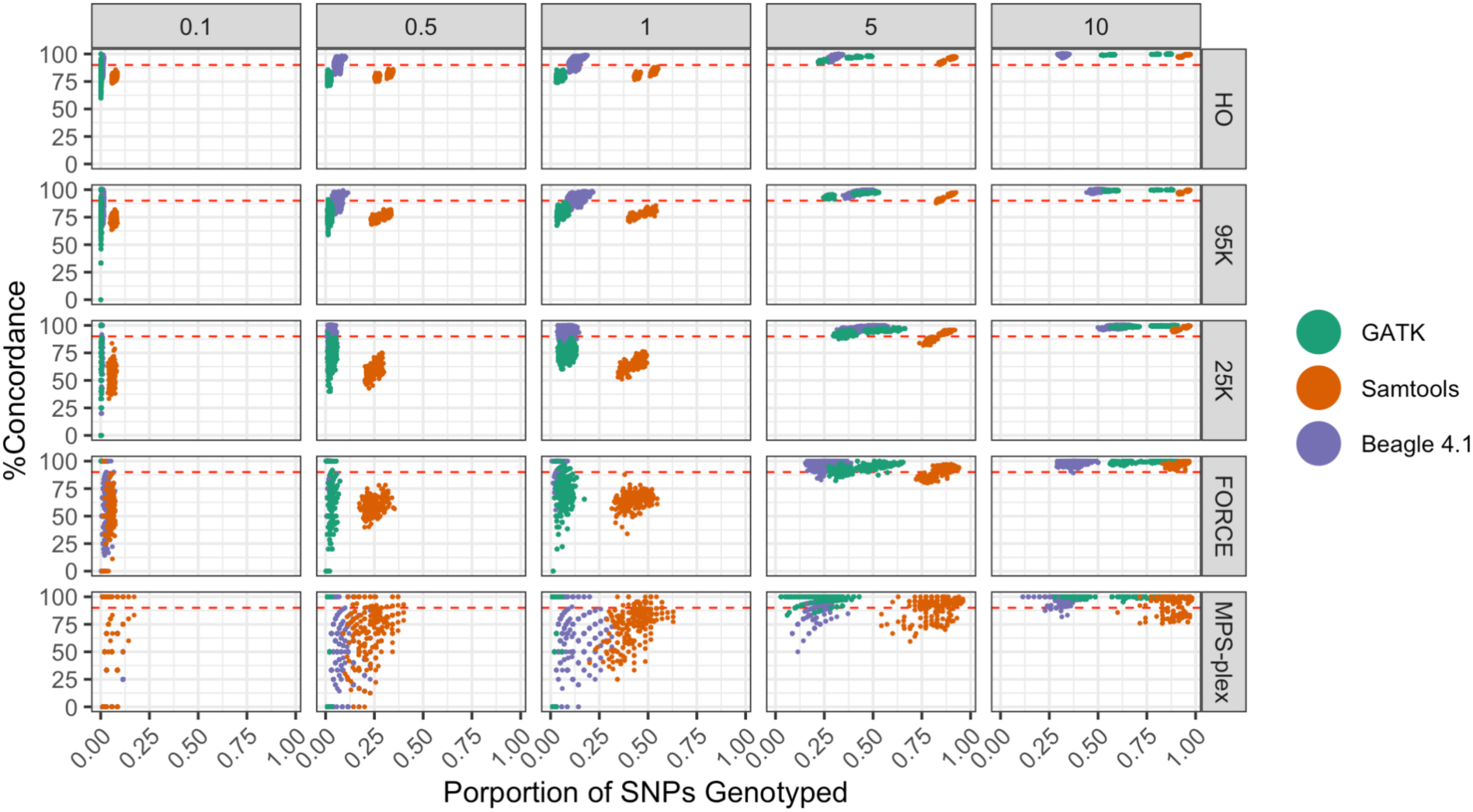
Concordance of genotypes compared to proportion of SNPs genotyped from different SNP panels across genotype refinement methods. Each coverage, genotyper, and SNP panel combination is represented by simulated sequencing data from 40 individuals. Genotype refinements were filtered for 99% genotype likelihood. For refinement with GATK and Beagle 4.1, data represents all four tested population reference panels. All DNA qualities and population reference panels are included. The red dashed line denotes 90% concordance.

Similar to the observations of genotype concordance pre-refinement, calling heterozygous sites below 5X coverage depth resulted in increased error rates (**Figure S16**).

To examine the impact of genetic similarity between the individual of interest and the population reference panel used for genotype refinement accuracy, we calculated *FST* between each individual genome and the respective population reference panel used for analysis with Beagle 4.1 and GATK (**Methods 2.2**). Given the impact of SNP density on the effectiveness of genotype refinement, we restricted this analysis to the three largest SNP panels. We find that for coverages less than 5X there is a negative correlation between concordance and genetic distance (Pearson correlation test: correlations - 0.41 to -0.06, max p-value of 3.24E-2 for 26 of 27 tests; **Figure S17-19**). However, when a population reference panel that represents all 5 super populations in the 1000 genomes dataset is used this negative correlation was decreased and no longer significant in almost all datasets with at least 0.5X coverage (Pearson correlation test: correlations -.21 to 0.04, p-values > 5E-2 for 29 of 36 tests; **Figure S20-22**). Across methods, the ATLAS genotyper paired with Beagle 4.1 consistently results in higher concordance rates, though results in a lower number of genotyped SNPs (**Figures 5 and S14**).

SAMtools consistently retains the most SNPs post-refinement, in particular for more degraded DNA, albeit with lower concordance rates than the ATLAS + Beagle 4.1 combination. This observation may indicate that refinement methods utilizing haplotype information require more SNP density than the SNP panels included in this testing performed here.

### 3.5 Imputation

Imputation is commonly used in genetic studies to fill in missing genotypes based on haplotype information from population reference panels^52^. We tested two commonly used imputation methods, Beagle 5.4^43^ and GLIMPSE2^44,45^. As SNP density impacts imputation efficiency, we only used the larger three SNP panels (25K, 95K, and Human Origins) and then tested the performance for varying coverages between 0.5-10X. We used the output of genotype refinement as the input for imputation. The ATLAS and Beagle 4.1 was selected as it resulted in the highest concordances for lower coverages. However, due to the extreme decrease in total number of SNPs post-genotype refinement we also tested SAMtools paired with GLIMPSE2. As with genotype refinement we used a filter of 99% genotype probability to test genotype concordance.

For the largest SNP panel (the Human Origins panel), all parameters tested resulted in greater than 90% concordance when using a population reference panel representative of the genetic diversity of the 1000 genomes dataset (**Figure 6**) with outliers dipping below 90% when using other population reference panels (**Figure S23**). As expected, the concordance rates fall when using smaller SNP panels and individuals with lower coverages, likely due to the impact of decreased SNP density (median decrease in percent concordance for <5X coverage 2% and 8% for 95K and 25K panels respectively). We find that GLIMPSE2 consistently resulted in higher concordances (**Figure 6)** for the two largest SNP panels, though Beagle 5.4 retained the most SNPs (**Figures S24-25**) (Median concordance of 96% and 97% with median SNP panel coverage proportions of 0.49 and 0.40 for Beagle 5.4 and GLIMPSE2 respectively). The increased concordance in high coverages and larger SNP panels may be due to the larger number of SNPs retained post-refinement (**Figures S24-25**). SAMtools paired with GLIMPSE2 did result in lower concordances for coverages below 5X for the smallest SNP panel tested (25K panel). Post genotype refinement SAMtools also resulted in lower concordances (**Figure 5**), which may in turn further lead to reduced concordance after imputation. The increase in the number of genotyped SNPs was greatest for coverages <5X (**Figure S26**), though remained lower than the numbers of SNPs originally genotyped (**Figure 7**). Similar to the observations with genotype refinement, utilizing a population reference panel containing individuals from all super populations in the 1000 genomes dataset resulted in increased concordance across the range of observed *FST* values, regardless of matching the individuals to the closest genetically similar population (**Figures S27- S28**). However, for lower coverages, reduced concordance correlates with distance from the population reference panel (**Figure S30**). Comparing the performance by MAF, we find most common variants are well imputed though the concordance is lower for heterozygous rare variants (**Figure S29**). Using the population reference panel representative of all super populations in the 1000 genomes dataset increases the concordance for some MAF bins, but does not eliminate this trend for rare variants (**Figures S30-31**). It has been demonstrated in other studies that a diverse population reference panel supplemented with population-specific samples can improve concordance rates for rare variants^16^.

**Figure 6.**
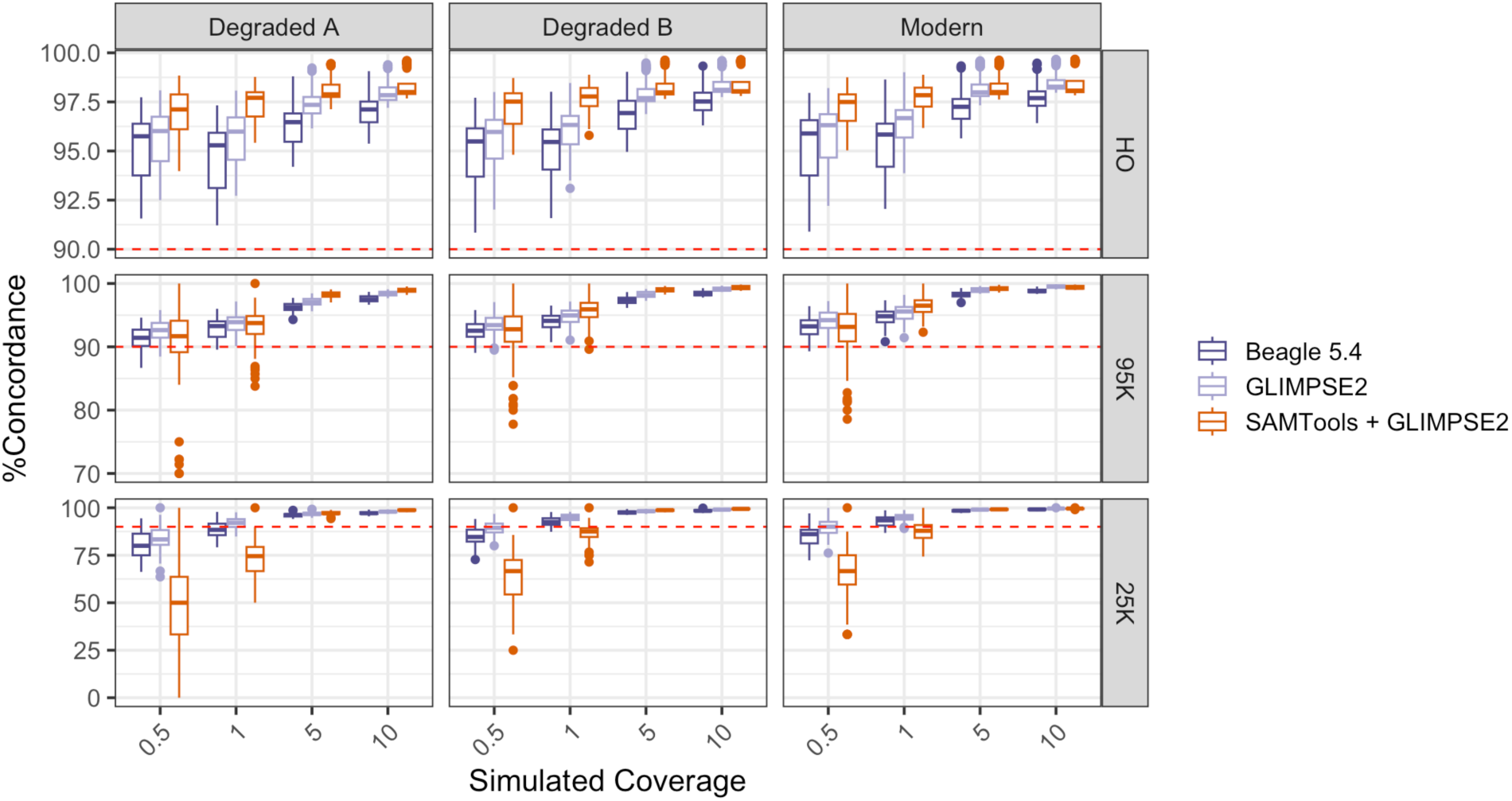
Concordance of genotypes for different workflows of genotyping and imputation methods using a population reference panel representative of the 1000 genomes dataset. Genotype likelihood thresholds of 99% were used for refinement and imputation. The Tukey box and whisker plots show the lower and upper quartiles boxed with the whiskers representing 1.5 times the interquartile range. Dots denote any outliers. Beagle 5.4: Atlas genotyping + Beagle 4.1 refinement + Beagle 5.4 imputation; GLIMPSE2: Atlas genotyping + Beagle 4.1 refinement +GLIMPSE2. Note that the y-axis is different for each SNP panel. The red dashed line denotes 90% concordance.

**Figure 7.**
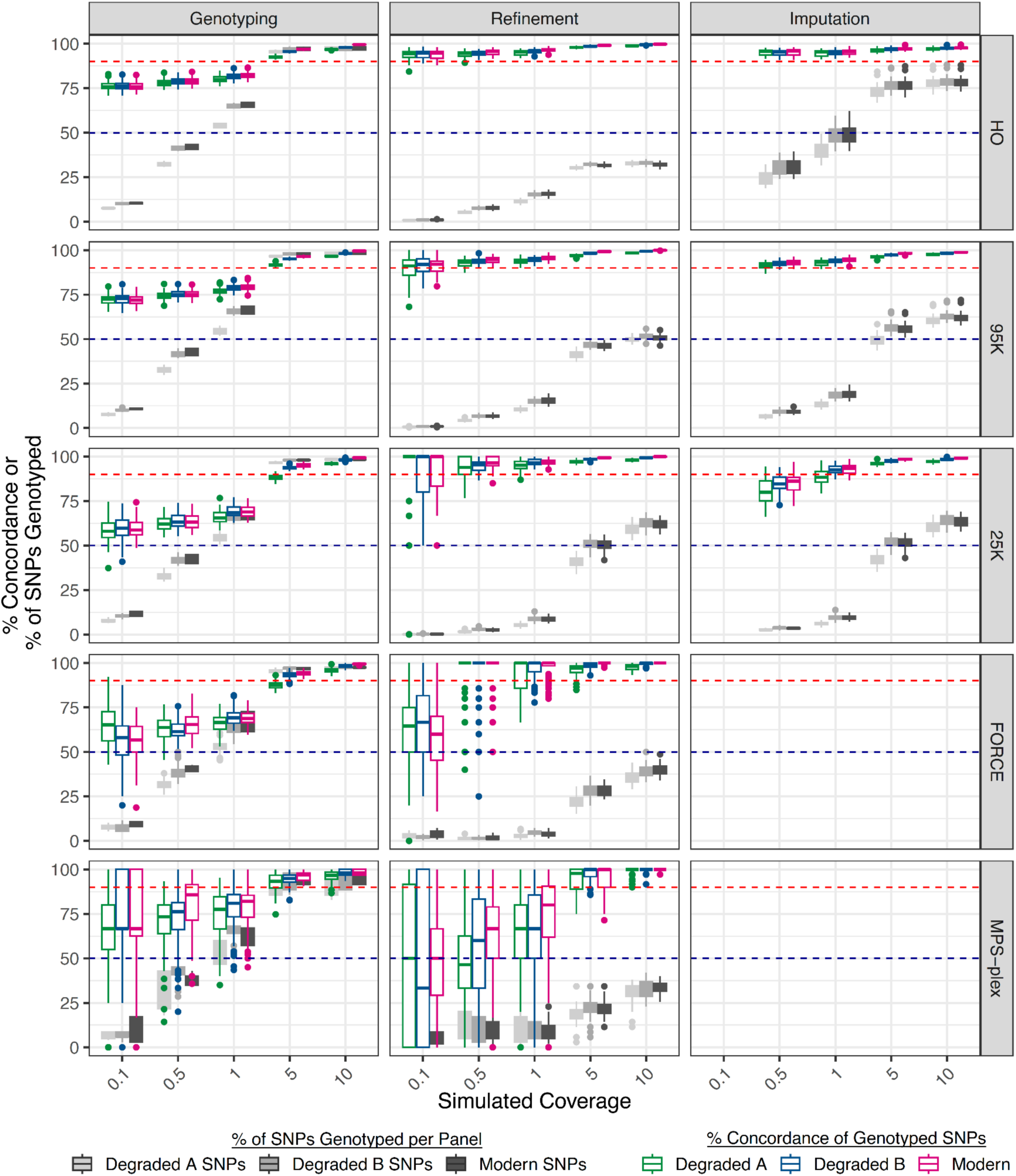
Concordance of Genotyped SNPs and breadth of Genotyped SNPs per SNP panel through different workflow steps. ATLAS genotyping followed by Beagle 4.1 refinement and Beagle 5.4 imputation using the 1000 genomes reference panel. Grey box plots represent % of SNPs in each SNP panel that were genotyped. Each coverage, DNA quality genotyper, and SNP panel combination is represented by simulated sequencing data from 40 individuals. Genotypes post-refinement and imputation were filtered for at least 99% genotype probability. The Tukey box and whisker plots show the lower and upper quartiles boxed with the whiskers representing 1.5 times the interquartile range. Dots denote any outliers. The red dashed line denotes 90% concordance, and the dark blue dashed line denotes 50%.

These studies also maintain the trend that larger SNP panels result in higher imputation accuracies, emphasizing that SNP density can have an even larger impact than sequencing depth for NGS data.

## 4. Discussion

In this study, we utilize simulations to understand the limitations of the computational analysis of genomic data for forensic applications using degraded DNA. We find leveraging some of the innovations in analyzing degraded data in ancient DNA research has significant impact on sequence mapping and filtering and increases the retention of degraded reads and higher genotype concordance rates, underscoring the relevance of using ancient DNA methods for forensic analysis. Indeed, this is already being recognized in forensics as exemplified by the integration of ATLAS into Parabon’s analytical pipeline^29^.

We find that reduced sequencing depth and increased degradation both result in decreased concordance rates across genotyping methods, with increased variance in genotype accuracy for smaller SNP panels (**Figure 7**). Genotype refinement methods that leverage population reference panels result in higher accuracies but require a large number of SNPs (typically thousands per chromosome) to be applicable. In turn, imputation is able to recover a portion of this loss while maintaining high accuracies, but still results in less complete DNA profiles than observed pre-refinement and imputation for higher coverages (**Figure 7**). These results emphasize the importance of SNP density for fully leveraging the power of genotype refinement and imputation methods that rely on population reference panels. For this reason, we do not recommend genotype refinement or imputation for data with coverages of 10X and greater, which in our evaluations resulted in >85% concordance while covering at least 80% of SNPs in the tri-allelic SNP panel and >90% concordance across at least 95% of SNPs in the other four SNP panels after the initial genotyping step (**Figure 7**). For coverages <5X of the two largest tested SNP panels (95K and HO), refinement and imputation result in concordances of at least 85% even down to 0.1X when using a population reference panel such as the full 1000 genomes dataset (**Figure 7**).

Traditionally, in the validation of new methods in forensic genetics, parameters such as DNA quantity and sequencing depth are key components for understanding methodological limitations and establishing reliable workflows^53^. In this study, we show that there may be reduced accuracy for imputing rare variants at low coverages. We also show that genotype concordance is impacted by heterozygosity and genetic similarity of the individual to the population reference panels used. As heterozygosity varies across worldwide populations and also availability of reference data differs by population groups, this can lead to differences in genotype accuracy across ancestry groups. For this reason, we suggest that guidelines for the validation of probabilistic genotyping, refinement, and imputation include testing on genotypes from individuals of a range of genetic ancestries. The use of continuous population descriptors as well performing evaluations across minor allele frequencies in these tests will make them more widely applicable. We suggest heterozygosity and *FST* as only starting points for evaluating the relationship between genetic ancestry and accuracy of methods analyzing sequencing data. This said, determining which modern population an individual has the greatest genetic similarity to, can still be informative in some contexts. While we find that using a reference database reflective of the full 1000 genomes dataset results in higher concordances for our SNP panels which primarily contain common variants, it has been shown in other studies that for rare variants using the 1000 genomes dataset combined with additional individuals that have low *FST* to the individual of interest increases accuracy even more^16^. Therefore, laboratories that may have their own internal reference databases that are tailored to relevant genetic ancestries may observe increased accuracy by combining them with the publicly available data.

A consistent trend for genotype refinement and imputation was a decrease of concordance with not only depth of coverage, but also size of the SNP panel. Historically, the forensic field has favored analyses that target specific loci due to cost, efficacy, and privacy concerns^47^. In the United States, CODIS STRs were selected in part due their being medically uninformative^54,55^, although this has since been called into question^56^. There are currently multiple NGS kits or SNP panels available for forensic analyses (94 - 9,959 SNPs ^48^), but most target relatively few SNPs compared to SNP panels in other fields (widely used commercial SNP arrays for medical and population genetic studies contain ∼700,000–3.7 millions SNPs^57–59^). Low coverage whole genome sequencing combined with refinement and imputation has become increasingly popular in place of high coverage sequencing in genomic studies to decrease sequencing costs. This raises the question if instead of developing new, smaller SNP panels, the forensic community should focus on larger SNP panels for applications with degraded DNA. Tests identifying the minimum number and spacing of SNPs needed for specific forensic analyses will also aid in decisions around ideal SNP panel sizes for degraded DNA. The issue of how many SNPs are needed to perform downstream forensic analyses also prompts the question as to what is the gold standard for genotype concordance. The threshold for concordance will also inform how to make decisions around the tradeoffs of decreased accuracy of smaller SNP panels with lower quality data and privacy concerns of larger SNP panels. Conventionally, 100% concordance has been considered the requirement for forensic analysis. However, as we see in our simulations, even at 10X coverage 0-13.3% error rates were observed. Incorporating these error rates into a likelihood ratio^60^ or propagating the uncertainty forward while calculating likelihood ratios directly from sequencing data^12^ will expand the amount and improve the reliability of data that could be analyzed in forensic casework.

## 5. Conclusion

In this study we provide a framework for testing genome mapping, genotyping, refinement, and imputation methods via simulation. Our tests show that sequencing depth impacts accuracy of genotyping. We also show that genetic ancestry (heterozygosity or genetic relationship to population reference panels used) impacts observed accuracy and therefore the importance that any validations of these methods should include DNA profiles reflective of a range of genetic ancestries. We also show that moving away from socially-determined or continental population groupings or descriptors in these evaluations will aid in creating more widely applicable guidelines that are reflective of the continuity of human genetic diversity. Furthermore, our tests reveal that SNP panels with low SNP density limit the power of genotype refinement and imputation methods. While genotype refinement and imputation do increase concordance rates for lower coverages, these steps still come with the trade-off of severely reducing the breadth of coverage across the genome. In these scenarios, it is necessary to consider the minimum number of SNPs needed to achieve the required degree of genotyping accuracy to be informative for the downstream analyses. In sum, we provide benchmarks for using next generation sequencing data from degraded DNA from diverse populations for forensics casework.

## Declaration of Competing Interest

The authors declare that they have no known competing financial interests or personal relationships that could have appeared to influence the work reported in this paper.

## Data Statement

The 1000 genomes high coverage data is publicly available and can be accessed here (https://ftp.1000genomes.ebi.ac.uk/vol1/ftp/data_collections/1000G_2504_high_coverage/). The TopMed data is available through dbGaP accession number phs001644.v3.p2. The workflow used for running simulations can be found at https://github.com/ezavala9/ForenDeg_Gt2Imp/ along with relevant scripts.

## Notes

### Competing Interest Statement

The authors have declared no competing interest.

