## Supplementary Material for "Benchmarking for genotyping and imputation using degraded DNA for forensic applications across diverse populations"

### 1. Simulation Parameters

In order to evaluate how degraded DNA may impact the accuracy of genotyping and imputation, we simulated data, varying three different parameters— sequencing depth, DNA quality, and SNP panels (Table S1). Degraded DNA leads to lower amounts of recovered genomic material and consequently lower sequence coverages. As the cost of sequencing scales with coverage depth, it is useful to understand the benefits of generating higher coverage data. Therefore, we evaluated five different sequencing depths (0.1, 0.5, 1, 5, and 10X). To evaluate impacts of DNA degradation, we simulated three different qualities of sequencing data: modern (sequences with no deamination and a uniform fragment size of 150 base pairs (bp)); degraded variation A (sequences with an average fragment size of ~40bp and terminal deamination of 20%); and degraded variation B (sequences with an average fragment size of ~80bp and terminal deamination of 10%). Moreover, as targeted sequencing is common in forensics due to both decreases in cost and a motivation to limit the generation of medically informative data[1], we used 5 SNP panels widely used in forensics applications including FORCE, MPS-plex, 25K and 95K panels as well as the Human Origins panel that is used in ancient DNA studies. After genotyping, VCFs were subset to the SNPs present in each of the respective SNP panels for subsequent analysis. We generated data for 40 individuals from the TopMed[2] and 1000 genomes phase 3[3] dataset.

**Table S1: Parameters for simulating sequencing data.**

| Parameter | Variables |
|---|---|
| Sequencing depth (X-fold) | 0.1, 1, 5, 10 |
| DNA Quality<br>(read length in base pairs (bp) and % of terminal deamination) | Degraded Variation A: ~40bp and 20% deamination<br>Degraded Variation B: ~80bp and 10% deamination<br>Modern: 150bp, 0% deamination |

| | |
|---|---|
| SNP Panel<br>(# SNPs excluding indels) | FORCE: 4,334 SNPs<br>MPS-plex: 1,209 SNPs<br>95K: 93,305 SNPs<br>25K: 24,914 SNPs<br>Human Origins: 600,706 SNPs |

### 2. Mapping and trimming

#### 2.1. Testing of mapping parameters

We performed the testing of genome mapping parameters using the published genotypes of 20 individuals from the 1000 genomes dataset[3]. NGSNGS[4] was used to simulate single end sequences of modern and degraded qualities (as described in Methods) with 0.1X coverage. We focused on SNPs in the FORCE and MPS-plex panels. The default *bwa* mapping parameters used were:

Default: bwa aln -t 16 {ref_genome} {fastq_file}

The ancient DNA parameters used were:

Ancient: bwa aln {ref_genome} -t 16 -n 0.01 -o 2 -l 16500 {fastq_file}

SAMtools v.1.8[5] was then used to remove sequences shorter than 35 bp or with mapping qualities less than 25.

To evaluate the impact of changing the mapping parameters, we examined: a) the ratio of all filtered reads mapped generated based on ancient or parameters (Figure S1A) and b) the difference in number of filtered mapped reads generated based on default or ancient parameters (Figure S1B). For modern DNA, we find the ratio is concordant with 1. However, for degraded samples, the ratio is significantly greater than 1 (paired Wilcoxon signed rank test, p-value = 7.8E-10), albeit it is only 0.6% higher at coverage of 0.1X but this difference translates to greater than 20,000 reads. Retaining the largest number of reads that can be critical especially for low coverage and degraded samples so we decided to use the ancient DNA parameters for subsequent analyses.

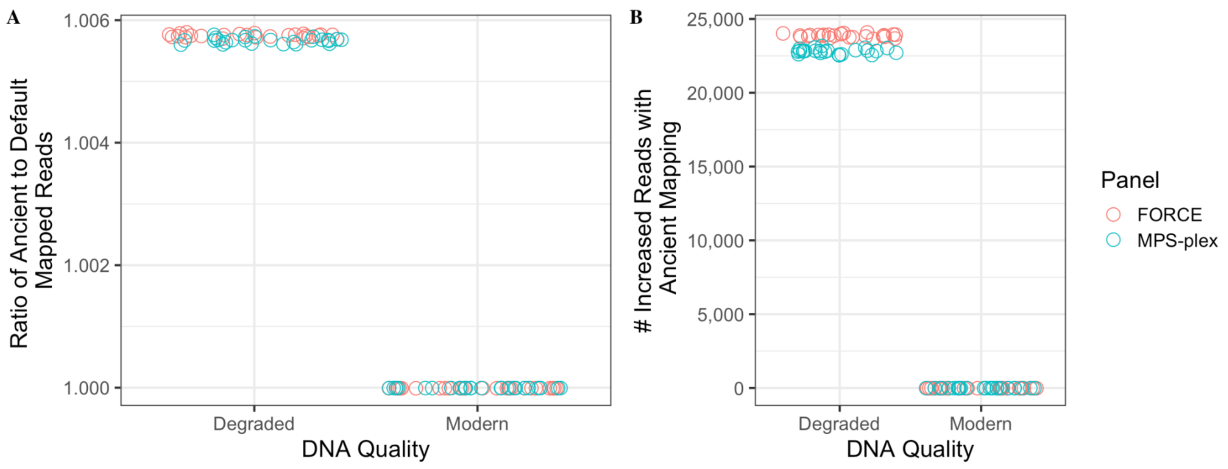


**Figure S1 Comparisons of the genome mapping parameters for whole genome sequences.** We show (A) ratio and (B) difference between the number of mapped reads using ancient or the bwa aln default parameters after filtering for length (≥35 bp) and mapping quality (25).

#### 2.2. Testing of mapping parameters for single chromosome simulations

Simulations to test the accuracy of genotyping and imputation were performed on chromosomes 1 and 14 (the largest chromosome and a mid-size chromosome) in order to decrease computing time. Therefore, we repeated the analysis in SI2.1 with simulating data only from chromosomes 1 and 14 (rather than whole genome sequences) and mapping to the human reference genome using "ancient" mapping settings. NGSNGS was used to perform simulations, focusing on two SNP panels (FORCE and MPS-plex) and varying data qualities (modern, degraded A, degraded B). We found that simulated modern sequences resulted in nearly zero mismapped reads. The degraded sequences had a higher proportion of mismapped reads than modern, though they were still under 0.01% (Figure S2). The mismapping rates mirror the patterns seen with simulated whole genome sequences (SI2.1; top two rows compared to bottom row) and hence for computational tractability, we focus on chromosomes 1 and 14 for the rest of the analyses.

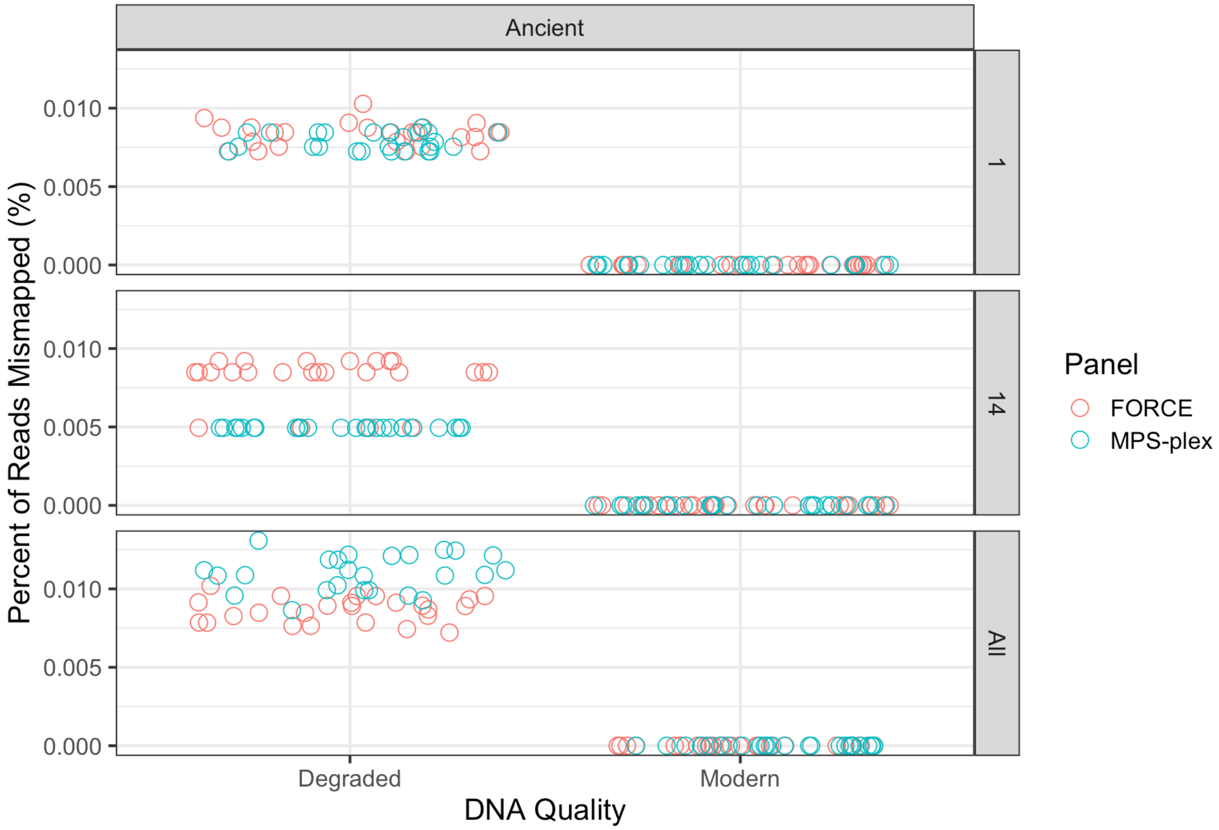


**Figure S2: Percent of simulated reads from chromosomes 1 and 14 that were mapped to incorrect regions of the human genome (hg38).** The first two rows represent simulated reads from a single chromosome. The third row represents data from section 2. 1, where reads were simulated across all chromosomes, subset to only reads that mapped to chromosomes 1 and 14.

#### 2.3. Trimming of terminal bases

In order to test the impact of trimming the terminal bases of read sequences, we simulated degraded DNA (variation A as described above) for varying coverages. We generated data for fifty individuals from the 1000 genomes dataset (Table S2). Trimming of the terminal bases was either performed before mapping (using the fastq file), post mapping and filtering (using the bam file), or not at all. We focused on removing the deaminated substitutions via trimming or masking based on the observed simulated deamination rates, which showed elevated substitution rates for the terminal 8 bases (Figure 2). Before mapping, one cannot determine the direction in which the read maps to the reference genome (forward or reverse) and hence for trimming at the fastq level, we removed the first and last eight bases of reads (a total of 16 bases). For trimming at the bam file level, we replaced C-to-T substitutions in the first 8bp on the 5' end and G-to-A substitutions from the first 8bp of the 3' end with an "N". This removed deamination from the bam file, but this approach does not reduce the read length (Figure S3). Mapping was performed as described above with bwa aln with ancient parameters. Genotyping was performed using ATLAS[6] following the settings described in Methods (Section 2.4) for a set of SNPs from chromosomes 1 and 14 in the FORCE panel. Concordance was then determined by comparing the resulting genotypes to the expected genotypes used for simulations from each dataset.

We find that there are marked differences between the three settings––NoTrim, fasTrim, bamTrim–– which are coverage dependent. Overall, each setting has increased genotype concordance with increased coverage. For 0.1X coverage there is no significant difference between the different settings (Figure S4, Table S3). Surprisingly trimming before mapping (fasTrim) resulted in a significantly lower consensus rate than the other two settings (bamTrim and NoTrim, min p-value = 5.8E-6) for sequencing depth of 1X to 5X. This may be caused by the decreased fragment sizes resulting from trimming of fastq file (Figure S3), leads to shorter overall fragment sizes, and in turn less sequences at all subsequent stages that is further exacerbated by filtering (35bp and mapping quality of 25). The impact of reduced number of sequences after filtering has less impact at higher coverages. As for bamTrim only transitions are masked, not all 8 terminal bases, these data are not impacted in the same way. It should be noted the modern sequences are longer and hence the extended trimming at the fastq level does not impact genotype concordance. As bamTrim outperforms other settings for all coverages (0.1-10X), we selected to perform trimming of sequences after mapping and filtering.

**Table S2. Sample identifiers used for simulations.** We simulated data for fifty individuals from the 1000 Genomes Project. We show the individual ID and super population descriptors as predefined by the 1000 Genomes Project. Bold IDs indicate individuals used subsequent testing of genotyping, refinement, and imputation methods.

| Individual | Super Population | Individual | Super Population | Individual | Super Population |
|---|---|---|---|---|---|
| **NA12778** | **European** | NA18954 | East Asian | NA19658 | American |
| **NA12812** | **European** | NA18959 | East Asian | NA19660 | American |
| **NA19717** | **American** | NA18961 | East Asian | NA19665 | American |
| **NA19729** | **American** | NA18528 | East Asian | NA19747 | American |

| | | | | | |
|---|---|---|---|---|---|
| **NA19700** | **African** | NA18530 | East Asian | NA21135 | South Asian |
| **NA19914** | **African** | NA18535 | East Asian | NA21142 | South Asian |
| **NA18942** | **East Asian** | NA18542 | East Asian | NA20891 | South Asian |
| **NA18947** | **East Asian** | NA18547 | East Asian | NA20896 | South Asian |
| **NA20904** | **South Asian** | NA10860 | European | NA20877 | South Asian |
| **NA21116** | **South Asian** | NA10865 | European | NA20911 | South Asian |
| NA12829 | European | NA11832 | European | NA21109 | South Asian |
| NA12843 | European | NA11882 | European | NA21111 | South Asian |
| NA12874 | European | NA11894 | European | | |
| NA19731 | American | NA18484 | African | | |
| NA19750 | American | NA18489 | African | | |
| NA19755 | American | NA18504 | African | | |
| NA19921 | African | NA18523 | African | | |
| NA20276 | African | NA18511 | African | | |
| NA19818 | African | NA19653 | American | | |

114

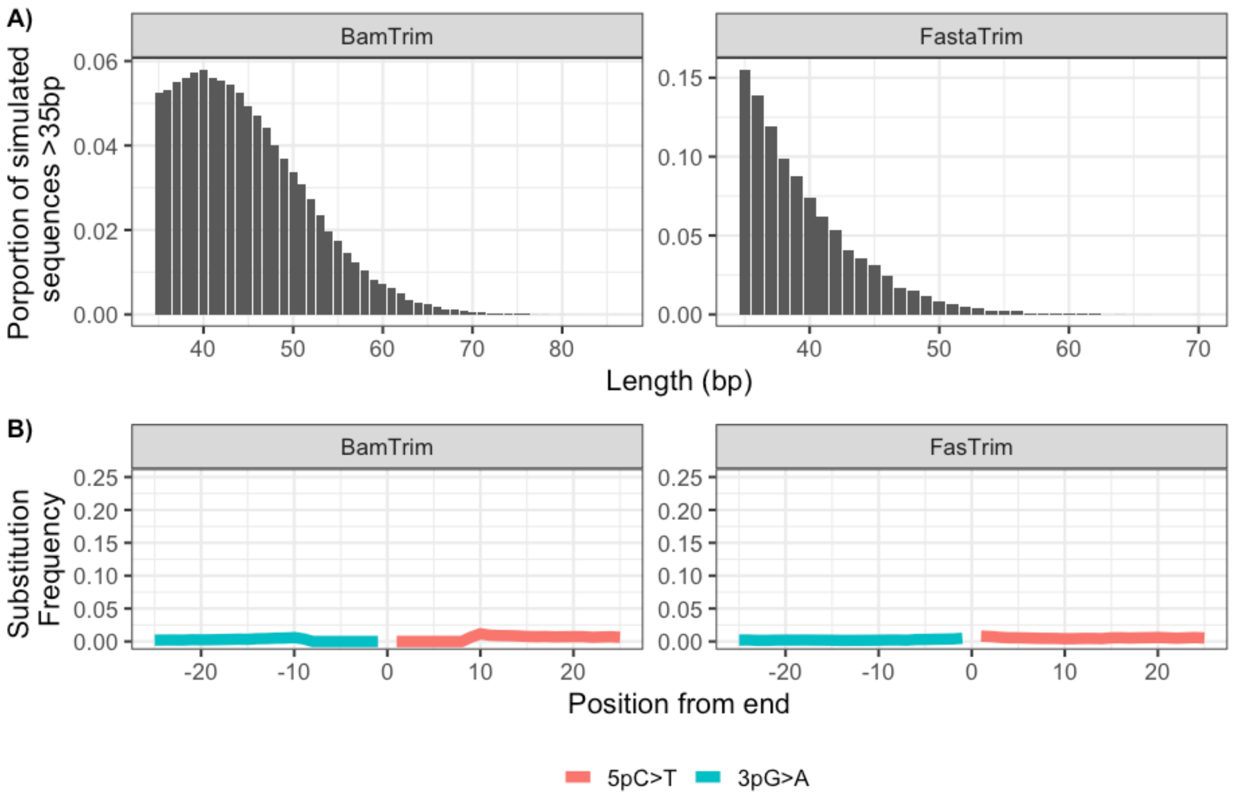


**Figure S3. Fragment size distributions and deamination patterns post trimming.** A) Fragment length distributions and B) substitution patterns used for simulated degraded variation A sequence data after trimming 8bps from the 5' and 3' ends of fastq sequences (FasTrim) or replacing putative deamination substitutions with N's in the first 8bps of 5' and 3' ends for mapped sequences (BamTrim).

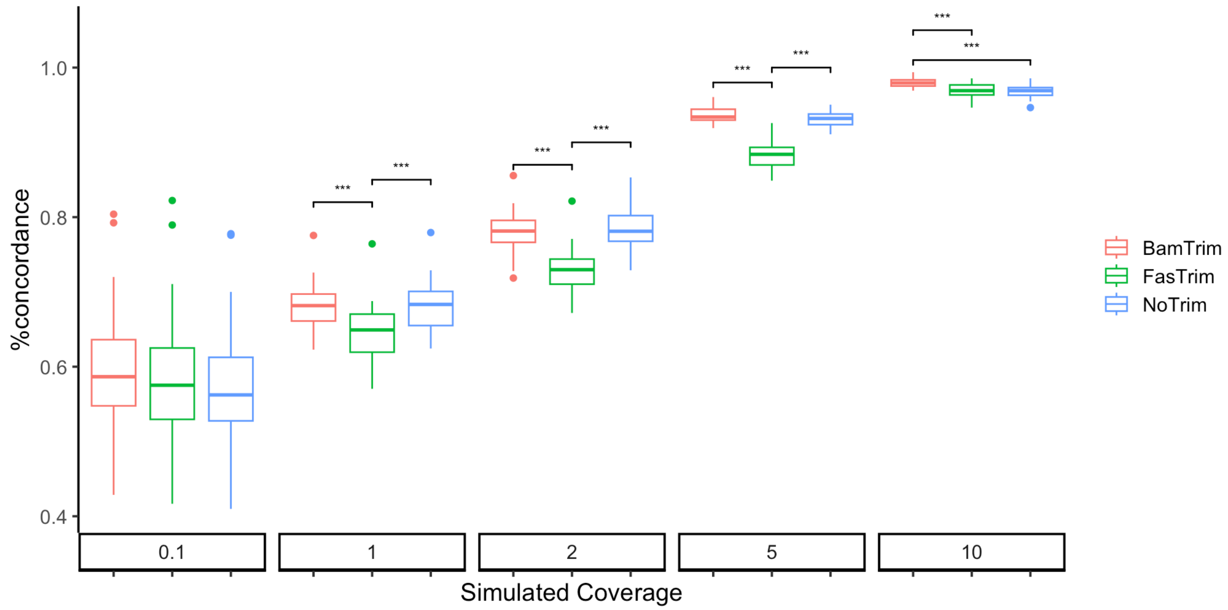


**Figure S4. Concordance of transition substitutions using different methods for removing deamination from simulated degraded A quality reads.** BamTrim - substitutions due to putative deamination were removed from first/last 8bps after mapping and filtering; FasTrim - 8bp the terminal ends were trimmed from the respective fastq file before mapping; NoTrim - no trimming was performed. Each coverage and trimming combination are represented by a subset of SNPs in 50 individuals from the 1000 genomes dataset. The Tukey box and whisker plots show the lower and upper quartiles boxed with the whiskers representing 1.5 times the interquartile range. Dots denote any outliers. Significant differences are noted with "***". See Table S3 for exact p-values.

**Table S3. Wilcoxon testing p-values with Bonferroni correction for multiple testing of genotype concordance for different trimming of simulated degraded A quality reads.** BamTrim - substitutions due to putative deamination were removed from first/last 8bps after mapping and filtering; FasTrim - 8bp the terminal ends were trimmed from the respective fastq file before mapping; NoTrim - no trimming was performed. Comparisons with a significant difference are noted in bold and red. Multiple testing correction performed with Bonferroni

| Sequence depth (range in #SNPs per individual) | BamTrim vs FasTrim | BamTrim vs NoTrim | NoTrim vs FasTrim |
|---|---|---|---|
| 0.1 (16-45) | 7.7E-1 | 3.1E-1 | 1 |
| 1 (120 -272) | 5.8E-6 | 1 | 3.1E-6 |
| 2 (196-389) | 1.4E-13 | 1 | 3.4E-14 |

| | | | |
|---|---|---|---|
| 5 (403-463) | 4.6E-17 | 7.7E-2 | 1.3E-16 |
| 10 (459-483) | 1.1E-8 | 5.5E-10 | 1 |

### 3. Genotype Concordance

We evaluated the impact of DNA quality on the accuracy of genotyping for different SNP panels for individuals from diverse populations by using three different genotypers commonly used in ancient DNA and forensic genetic analyses: ATLAS(Link et al., 2017), GATK(DePristo et al., 2011; McKenna et al., 2010; Van der Auwera & O'Connor, 2020), and SAMtools(Bonfield et al., 2021; Danecek et al., 2021). We simulated sequencing data as described in SI1 and Methods 2.1 from chromosomes 1 and 14 (the largest chromosome and a mid-size chromosome) in order to decrease computing time. The resulting sequences were mapped using "ancient" mapping settings, then filtered to keep higher quality reads (≥35 bp in length with mapping quality of ≥25) and any putative deamination on the terminal ends was masked with "N". The resulting reads were then used for genotyping. See Methods 2.5 for the settings used for each genotyper.

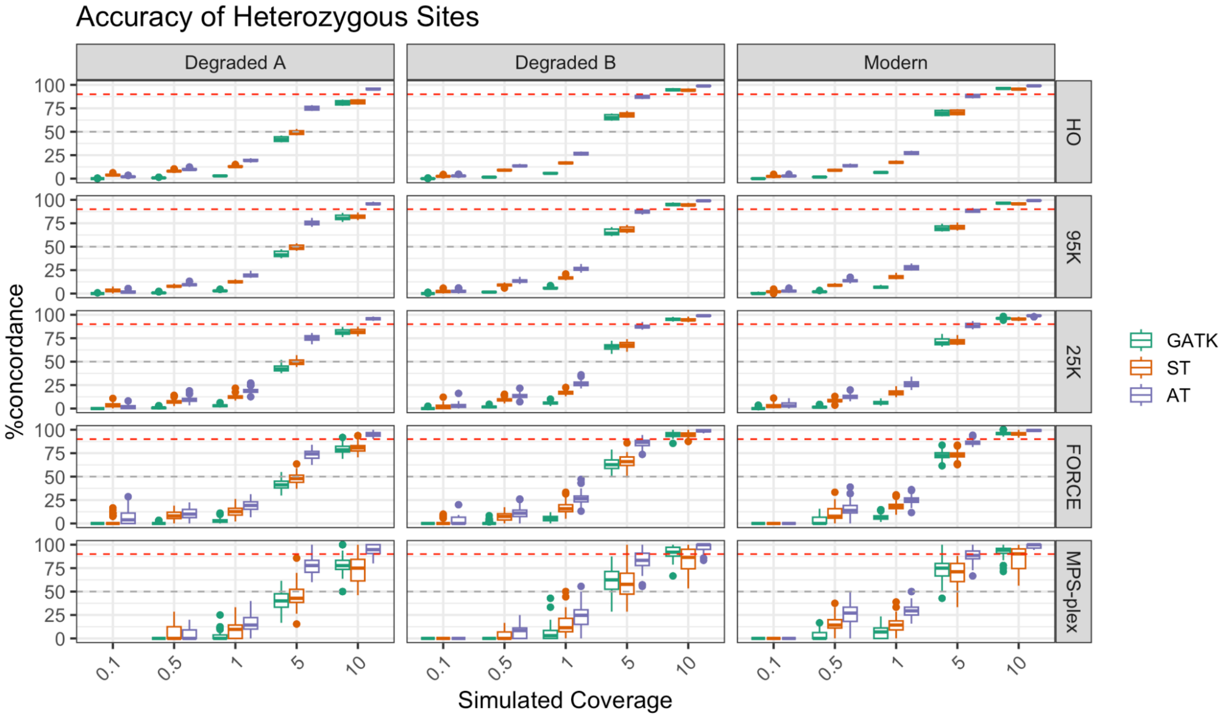

**Figure S5: Concordance across all heterozygous sites (0/1, 1/2, 0/2) per chromosome for each individual.** We require at least five heterozygous sites per individual per parameter combination. The red dotted line denotes 90% concordance and the gray dotted line 50%. Different genotypers are denoted by color (GATK = Genome Analysis Toolkit, ST= SAMtools and AT = Atlas). Each line represents a separate panel (HO = Human Origins). The Tukey box and whisker plots show the lower and upper quartiles boxed with the whiskers representing 1.5 times the interquartile range. Dots denote any outliers.

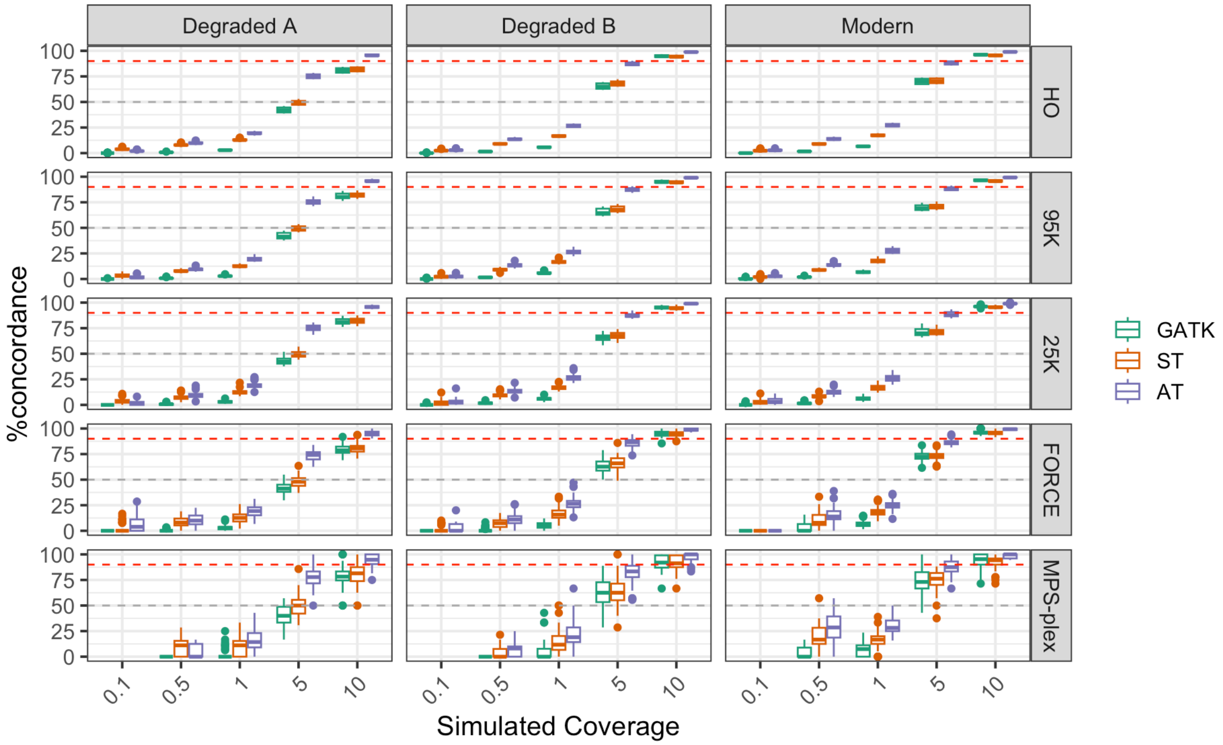


**Figure S6: Concordance across 0/1 heterozygous sites per chromosome for each individual.** We require at least five heterozygous sites in an individual per parameter combination. The red dotted line denotes 90% concordance and the gray dotted line 50%. Different genotype methods are denoted by color (GATK = Genome Analysis Toolkit, ST= SAMtools and AT = Atlas). Each line represents a separate panel (HO = Human Origins). The Tukey box and whisker plots show the lower and upper quartiles boxed with the whiskers representing 1.5 times the interquartile range. Dots denote any outliers.

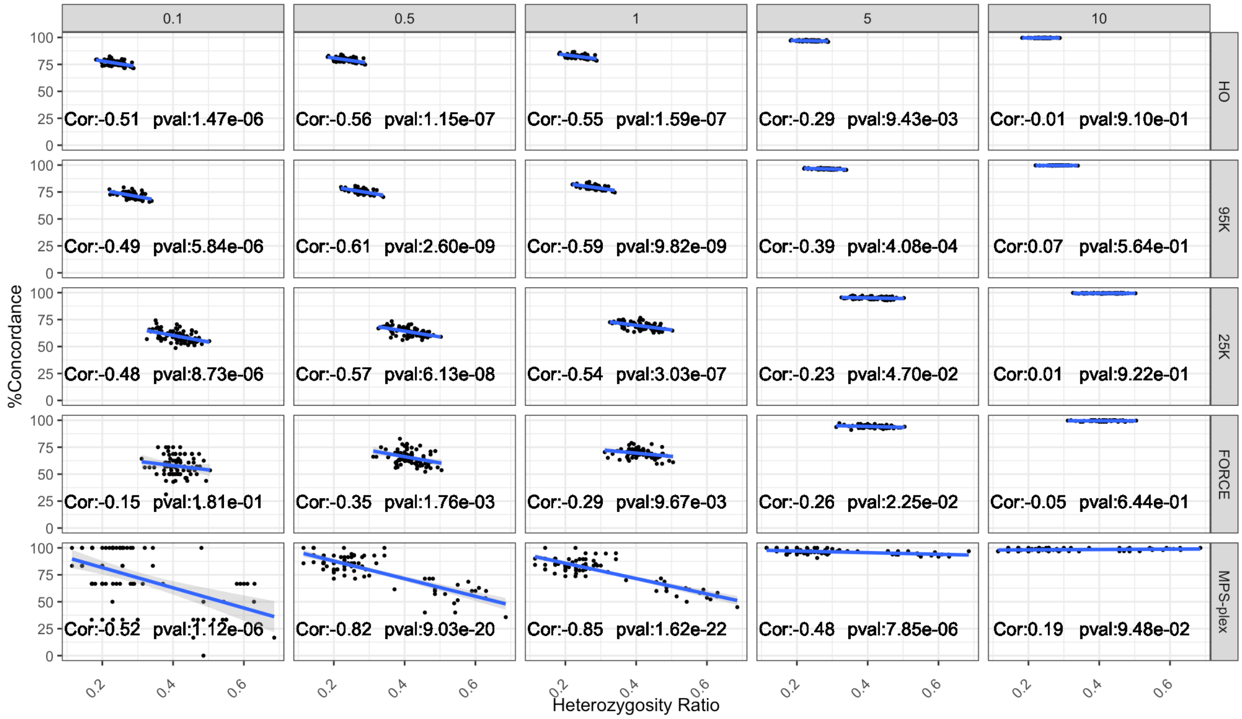


**Figure S7: Correlation between concordance and heterozygous ratio.** Heterozygous ratio was calculated as the proportion of sites per panel and chromosome (1 or 14) that have heterozygous genotypes divided by the number of SNPs on the SNP panel. Genotype concordance is based on genotyping using ATLAS. Correlation and p-values are based on Pearson's correlation test. The blue lines indicate the fit based on a linear model between the heterozygosity ratio and genotype concordance rates and the gray shaded area represents 95% confidence intervals.

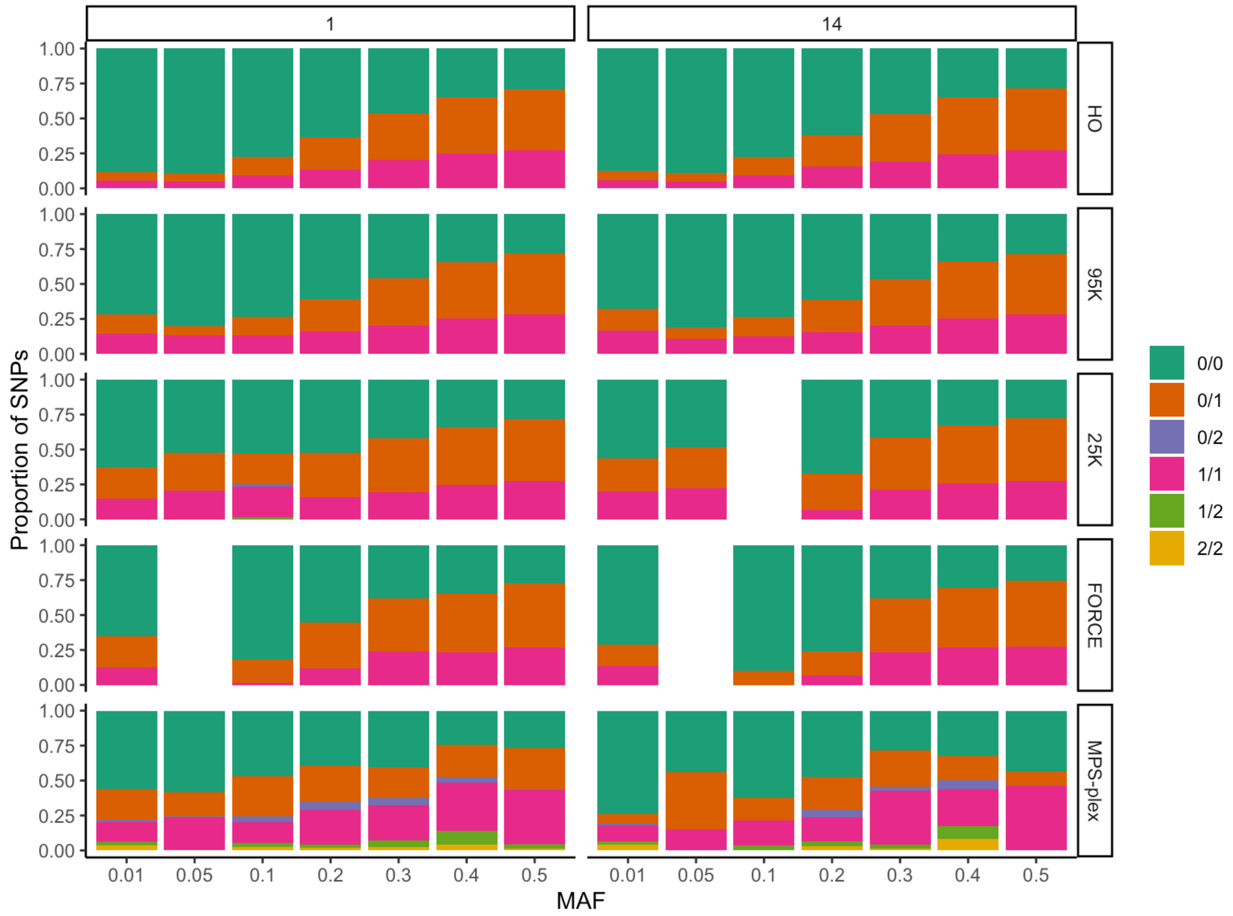


**Figure S8: Genotype frequency from simulated individuals across SNP panels and** minor allele frequency (MAFs). We represent the expected genotype data for chromosomes 1 and 14 from forty individuals used for simulations. Blanks indicate the respective panel did not cover genotypes with MAFs in the respective bin. MAFs are binned as follows: 0.01=0-0.01, 0.05=0.011-0.05; 0.1=0.051-0.1; 0.2=0.11-0.2; 0.3=0.21-0.3; 0.4=0.31-0.4; 0.5=0.41-0.5

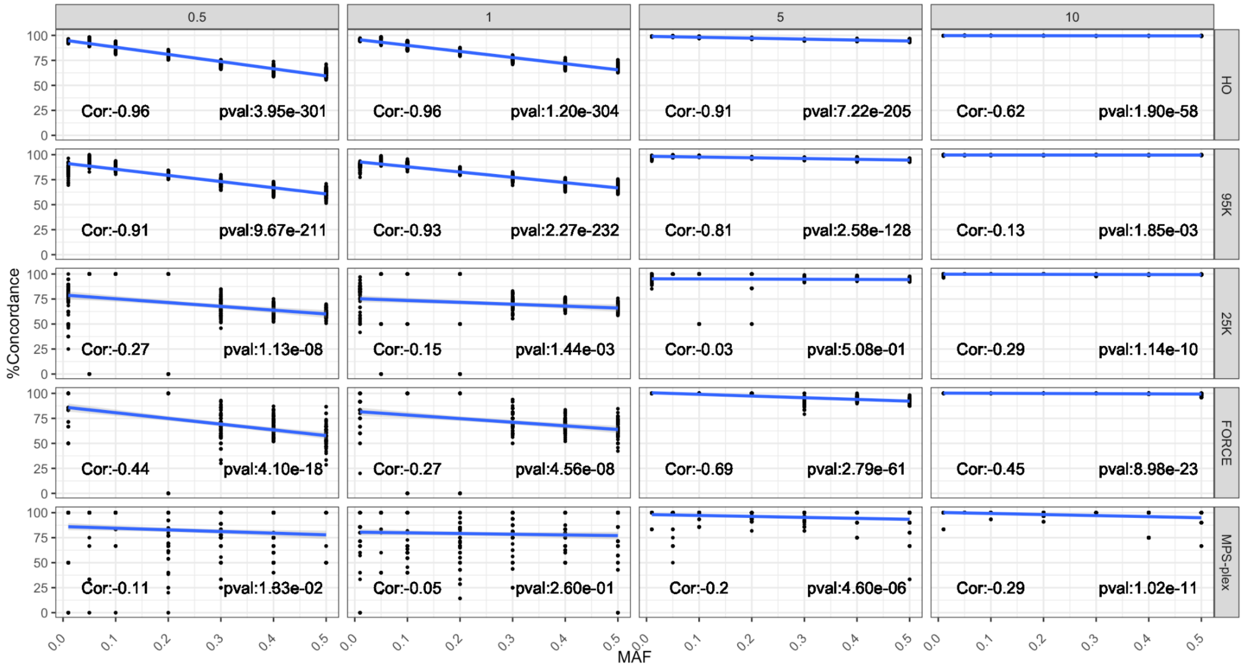


**Figure S9: Correlation between genotype concordance and minor allele frequencies (MAF).** Genotype concordance is based on the outputs from genotyping using ATLAS. Correlation and p-values are based on Pearson's correlation test. The blue lines are based on a linear model and the gray shaded area represents 95% confidence intervals. MAFs are as follows: 0.01=0-0.01, 0.05=0.011-0.05; 0.1=0.051-0.1; 0.2=0.11-0.2; 0.3=0.21-0.3; 0.4=0.31-0.4; 0.5=0.41-0.5

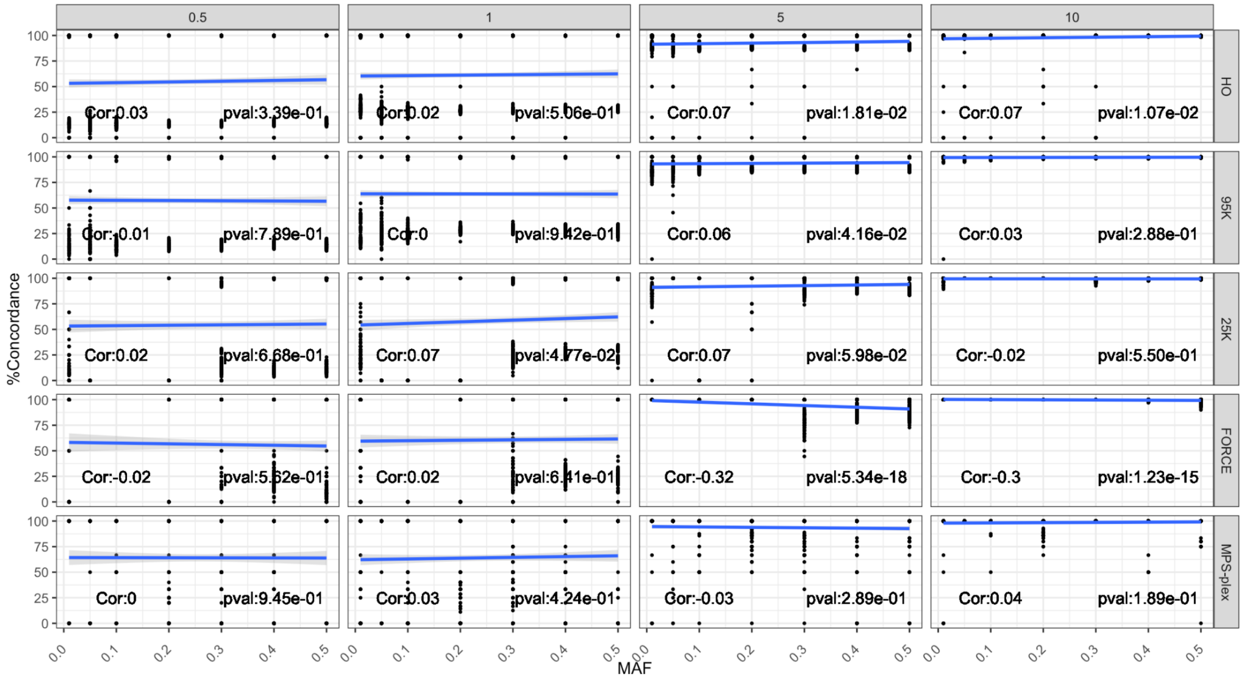


**Figure S10: Correlation between heterozygous genotype concordance and minor allele frequencies (MAF).** Genotype concordance is based on the outputs from genotyping using ATLAS.

Correlation and p-values are based on Pearson's correlation test. The blue lines are based on a linear model and the gray shaded area represents 95% confidence intervals. MAFs are as follows: 0.01=0-0.01, 0.05=0.011-0.05; 0.1=0.051-0.1; 0.2=0.11-0.2; 0.3=0.21-0.3; 0.4=0.31-0.4; 0.5=0.41-0.5

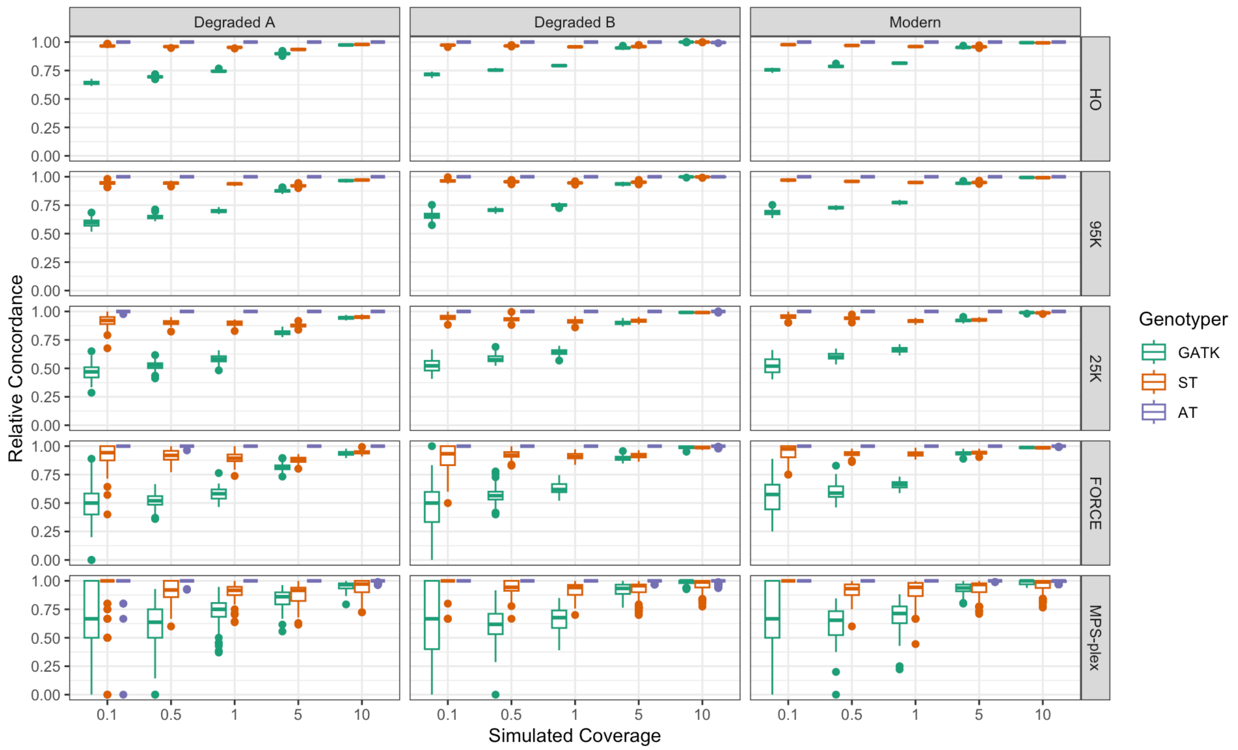


**Figure S11: Relative concordance compared to the genotyper with the most concordant SNPs for chromosomes 1 and 14 of all five SNP panels.** Each coverage, DNA quality, genotyper, and SNP panel combination is represented by simulated sequencing data from chromosomes 1 and 14 from 40 individuals. The Tukey box and whisker plots show the lower and upper quartiles boxed with the whiskers representing 1.5 times the interquartile range. Dots denote any outliers.

### 4. Genotype Refinement

Genotype refinement was performed on the outputs of each genotype method according to the recommendations for each respective method (**Methods 2.5**). For refinement of genotypes using ATLAS and GATK four different population reference panels were used, each consisting of 450 individuals. $F_{ST}$ was calculated to describe the genetic affinity between each simulated individual and the population reference panel (**Methods 2.2**).

**Table S4. Wilcoxon testing p-values with Bonferroni correction for multiple testing of change in genotype concordance after genotype refinement.** We assessed the performance for SNPs on chromosomes 1 and 14 for all three DNA quality types. Comparisons with a significant difference are noted in bold and red. Multiple testing correction performed with Bonferroni test. Genotypes were filtered for 99% genotype likelihood. For GATK and ATLAS the population reference panel of 450 individuals across all five super populations in 1000 genomes was used.

| Simulated sequence depth | Genotyper | SNP Panels (range in #SNPs per individual and step) | | | | |
|---|---|---|---|---|---|---|
| | | MPS-Plex | FORCE | 25K | 95K | HO |
| 0.1 | ATLAS | **3.2E-10 (1 - 8)** | 5.2E-2 (1 - 40) | **<2E-16 (1 - 245)** | **<2E-16 (13 - 825)** | **<2E-16 (126 - 5,212)** |
| | GATK | NA | **2.90E-14 (1 - 31)** | **<2E-16 (1 - 218)** | **<2E-16 (1 - 769)** | **<2E-16 (11 - 4,855)** |
| | SAMtools | 5.4E-1 (1 - 8) | **4.3E-2 (1 - 40)** | 1.8E-1 (31 - 246) | **<2E-16 (174 - 828)** | **<2E-16 (1,202 - 5,277)** |
| 0.5 | ATLAS | **1.2E-13 (1 - 50)** | **<2E-16 (1 - 149)** | **<2E-16 (9 - 889)** | **<2E-16 (112 - 3,251)** | **<2E-16 (949 - 21,333)** |
| | GATK | 8.8E-1 (1-46) | **<2E-16 (1 - 131)** | **<2E-16 (10 - 808)** | **<2E-16 (38 - 3,059)** | **<2E-16 (218 - 20,300)** |
| | SAMtools | **1.8E-4 (4 - 50)** | **6.4E-6 (26 - 150)** | **2.3E-7 (173 - 898)** | **<2E-16 (824 - 3,288)** | **<2E-16 (5,309 - 21,435)** |
| 1 | ATLAS | **6.8E-4 (1-70)** | **<2E-16 (2 -232)** | **<2E-16 (31 - 1,379)** | **<2E-16 (302 - 5,207)** | **<2E-16 (2,053 - 33,617)** |
| | GATK | **<2E-16 (1 - 68)** | **<2E-16 (2 - 216)** | **<2E-16 (35 - 1,300)** | **<2E-16 (109 - 4,909)** | **<2E-16 (671 - 32,462)** |
| | SAMtools | **1.6E-5 (10 - 70)** | **9.2E-9 (49 - 232)** | **4.9E-14 (295 - 1,393)** | **<2E-16 (1,403 - 5,259)** | **<2E-16 (8,971 - 33,717)** |
| 5 | ATLAS | **1.5E-8 (1 - 99)** | **<2E-16 (30 -340)** | **<2E-16 (307 - 2,051)** | **<2E-16 (1,301 - 7,763)** | **<2E-16 (6,042 - 50,647)** |

| | | | | | | |
|---|---|---|---|---|---|---|
| | GATK | <2E-16 (1 - 99) | <2E-16 (41 - 340) | <2E-16 (283 - 2,052) | <2E-16 (917 - 7,765) | <2E-16 (4,804 - 50,615) |
| | SAMtools | 2.40E-11 (19 - 99) | <2E-16 (111 - 340) | <2E-16 (627 - 2,052) | <2E-16 (2,809 - 7,771) | <2E-16 (17,072 - 50,657) |
| 10 | ATLAS | <2E-16 (4 - 99) | 2.5E-16 (47 - 340) | <2E-16 (458 - 2,056) | <2E-16 (1,575 - 7,795) | <2E-16 (6,018 - 50, 883) |
| | GATK | <2E-16 (11 -99) | <2E-16 (87 - 340) | <2E-16 (541 - 2,056) | <2E-16 (1,918 - 7,794) | <2E-16 (11,349 - 50,882) |
| | SAMtools | 6.5E-7 (21 - 99) | 4.2E-13 (125 - 340) | <2E-16 (750 - 2,056) | <2E-16 (3,088 - 7,795) | <2E-16 (18,585 - 50,883) |

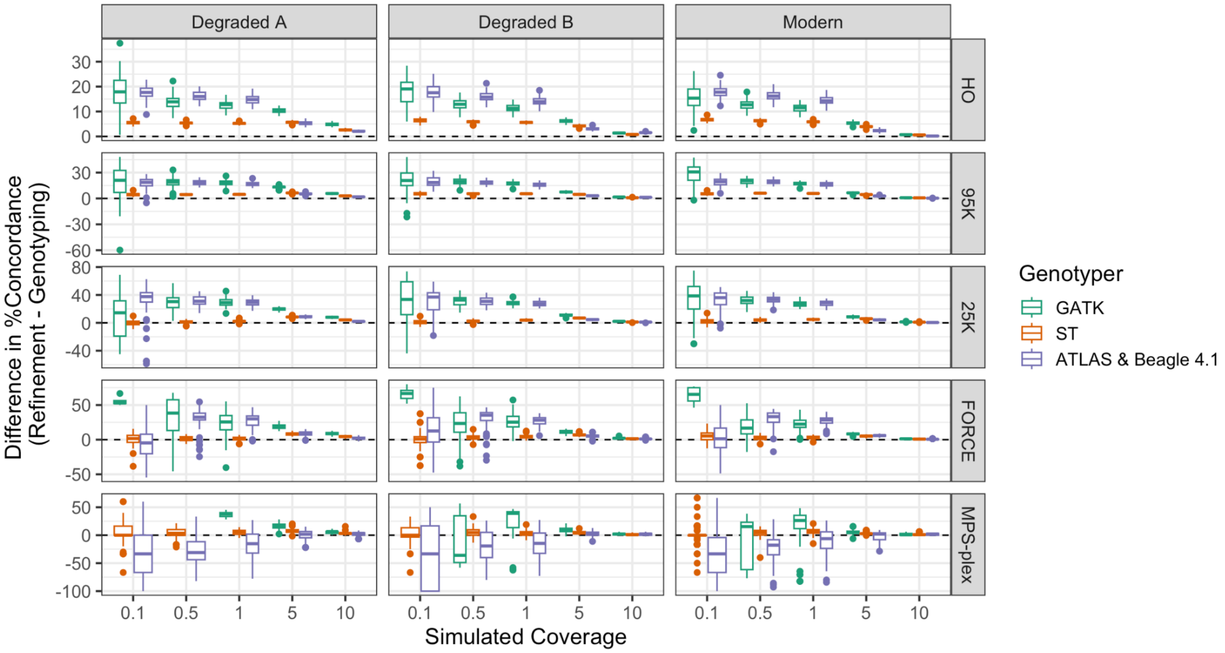


**Figure S12: The difference in percent genotype concordance between the refinement and genotyping steps.** Calculated by subtracting the percent concordance after genotyping from the concordance percentage after refinement for each combination of parameters. Note that the y-axis varies for each row. Genotype refinements were filtered for 99% genotype likelihood. For refinement with GATK and Beagle 4.1, the population reference panel of 450 individuals across all five super populations in the 1000 genomes dataset was used.

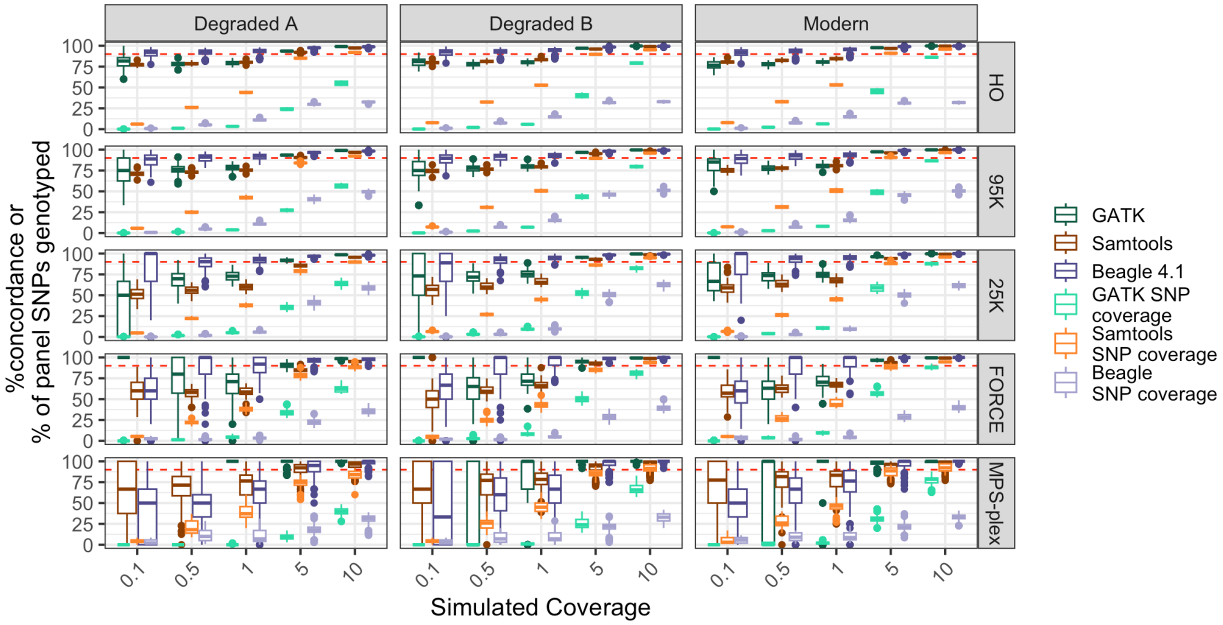


**Figure S13. Concordance of genotypes and breadth of SNP panel coverage for different genotyping refinement methods.** Each coverage, DNA quality genotyper, and SNP panel combination is represented by simulated sequencing data from 40 individuals. Genotype refinements were filtered for 99% genotype likelihood. For refinement with GATK and Beagle 4.1, data represents all four tested population reference panels. Breadth of SNP panel refers to the percent of SNPs for each respective SNP panel on chromosome 1 or 14 that were genotyped. The Tukey box and whisker plots show the lower and upper quartiles boxed with the whiskers representing 1.5 times the interquartile range. Dots denote any outliers. The red dashed line denotes 90% concordance or SNP breadth coverage.

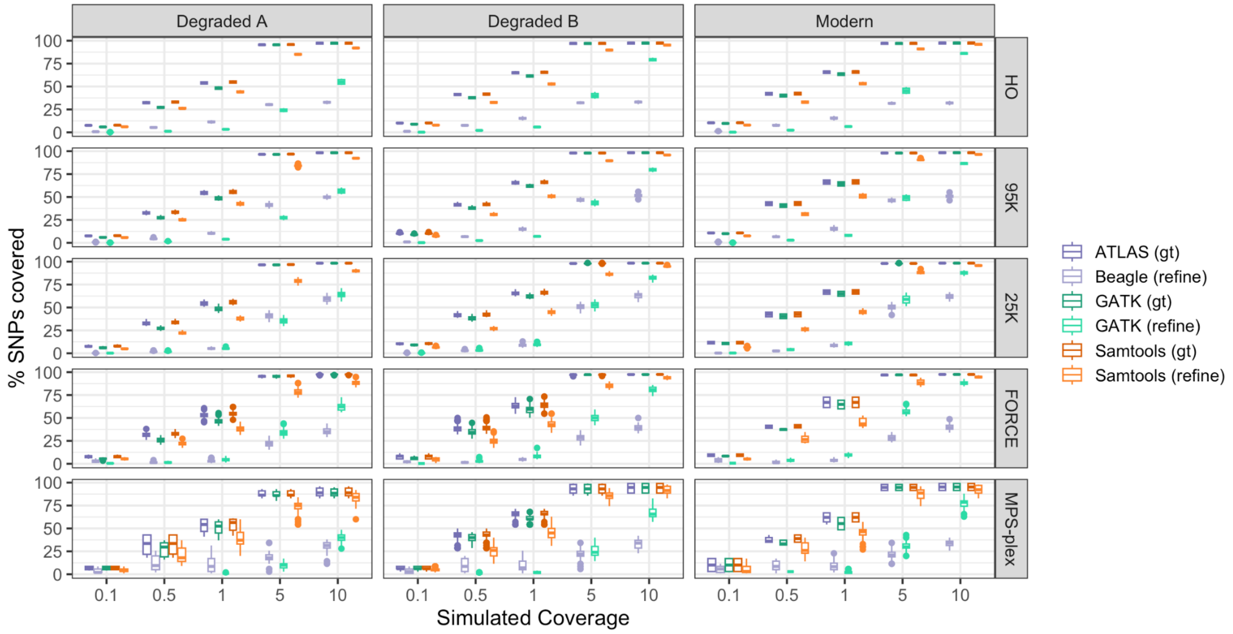


**Figure S14: Percent of SNPs genotyped in each panel before and after genotype refinement.** Genotype refinements were filtered for 99% genotype likelihood. For refinement with GATK and Beagle 4.1, the population reference panel of 450 individuals across all five super populations in the 1000 genomes dataset was used. Missing boxplots indicate no SNPs passed filters for that set of parameters. gt=Genotyping (before refinement), refine=post refinement. Percent SNPs was calculated based on the number of genotyped SNPs (passing filters) versus the expected number of SNPs per panel on either chromosome 1 or 14.

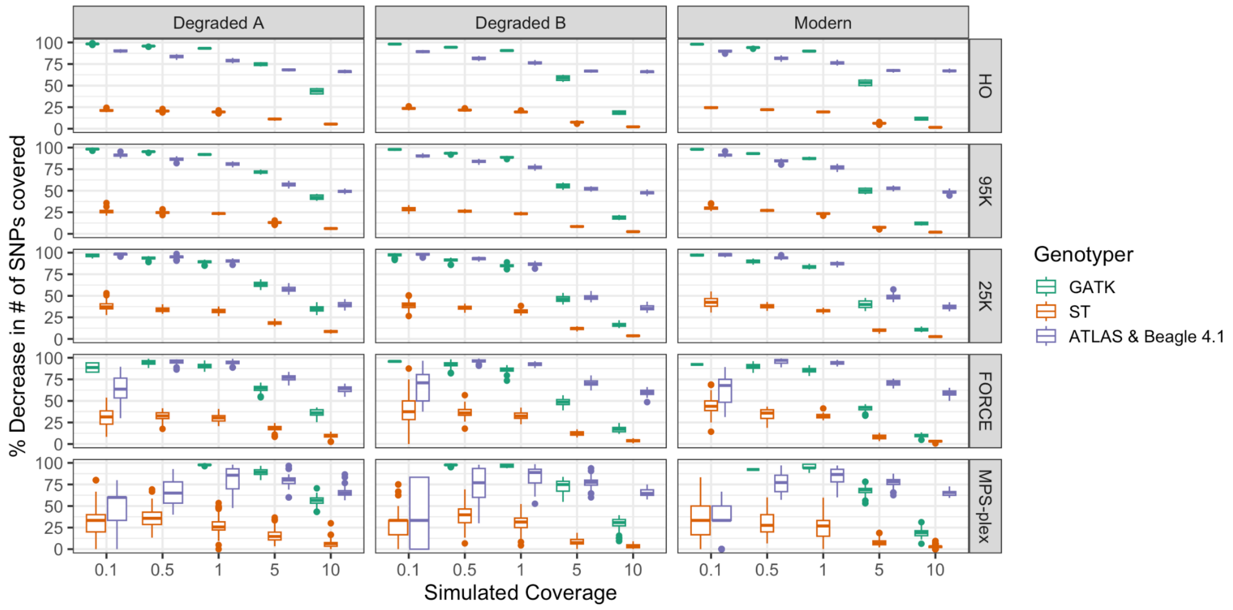

**Figure S15: Percent decrease in the number of SNPs covered after refinement.** We calculated the percent decrease as (#genotyped SNPs - #refined SNPS)/#genotyped SNPs. We filtered by removing genotypes with less than 99% genotype likelihood. For genotype refinement with GATK and Beagle 4.1, the population reference panel of 450 individuals across all five super populations in the 1000 genomes dataset was used.

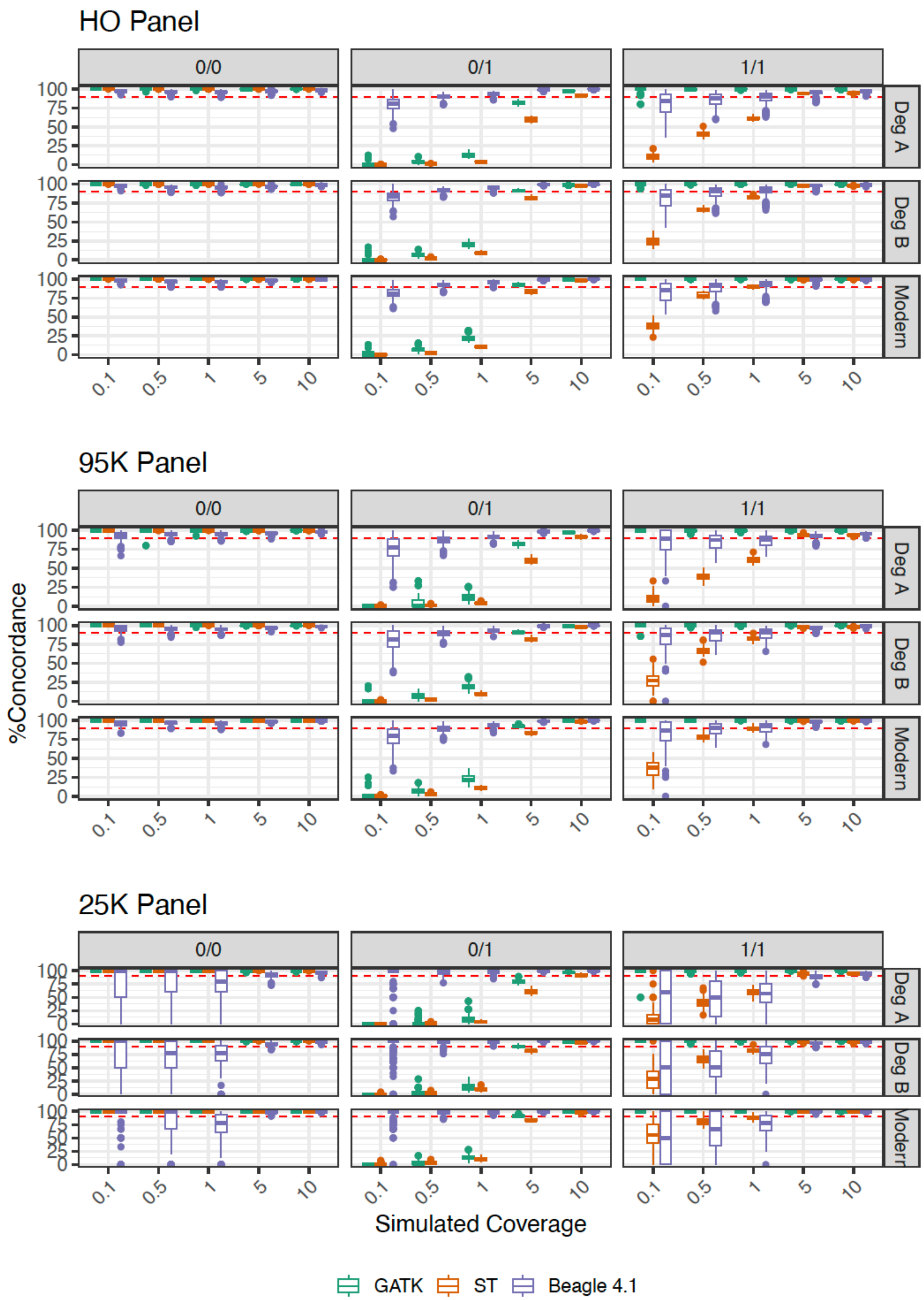
HO Panel
95K Panel
25K Panel
0/0
0/1
1/1
Deg A
Deg B
Modern
%Concordance
Simulated Coverage
0.1
0.5
1
5
10
0
25
50
75
100
GATK
ST
Beagle 4.1

260

**Figure S16: Genotype concordance across different genotypes.** We filtered genotypes removing those with less than 99% genotype likelihood. For genotype refinement with GATK and Beagle 4.1, the population reference panel of 450 individuals across all five super populations in the 1000 genomes dataset was used.

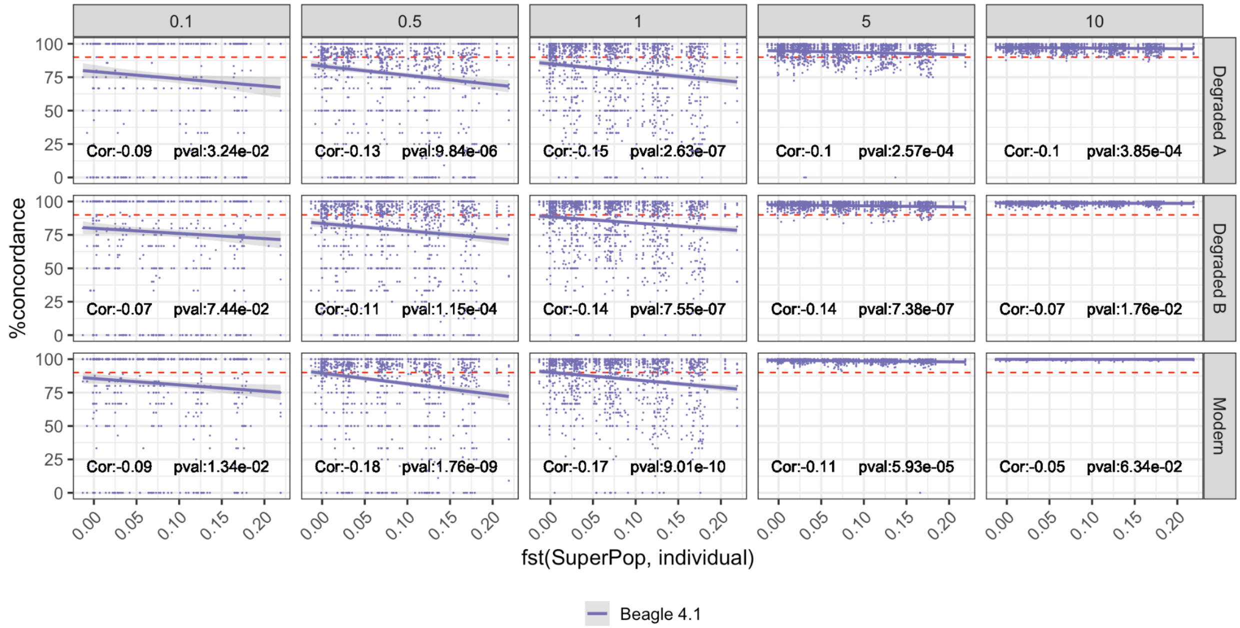


**Figure S17. Correlation between genotype concordance and genetic similarity between an individual and the population reference panel used for the 25K SNP panel.** We filtered genotypes removing those with less than99% genotype likelihood. Data for all tested population reference panels was included. Correlation and p-values were calculated using Pearson's correlation test. Higher $F_{ST}$ values indicate less genetic similarity.

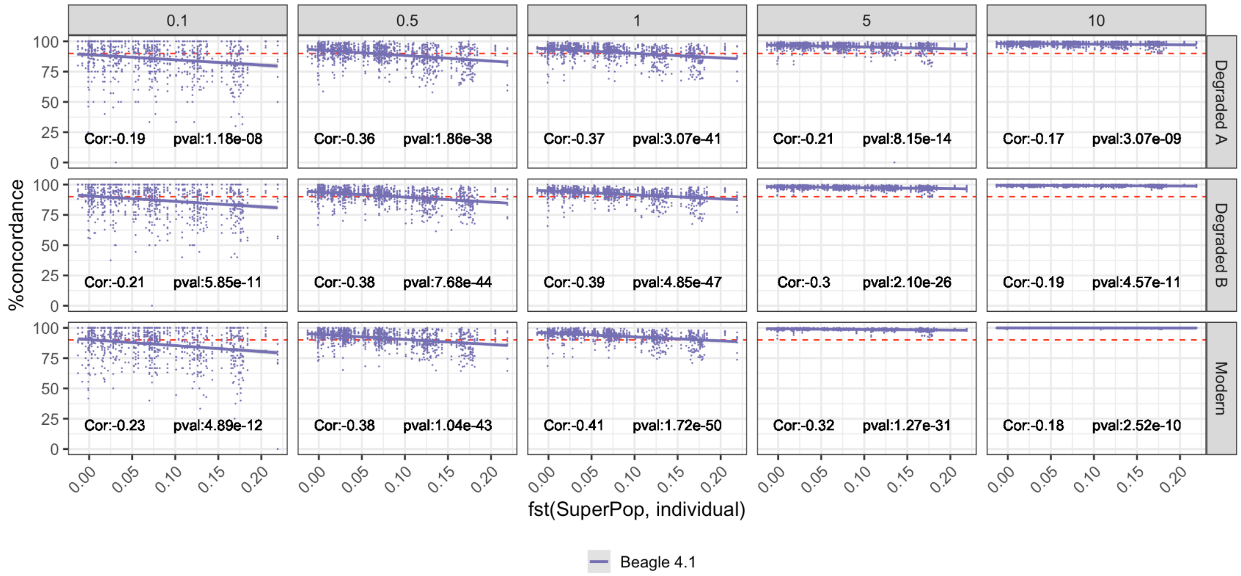

**Figure S18. Correlation between genotype concordance and genetic similarity between an individual and the population reference panel used for the 95K SNP panel.** Data for all tested population reference panels was included. Correlation and p-values were calculated using Pearson's correlation test. Higher $F_{ST}$ values indicate less genetic similarity.

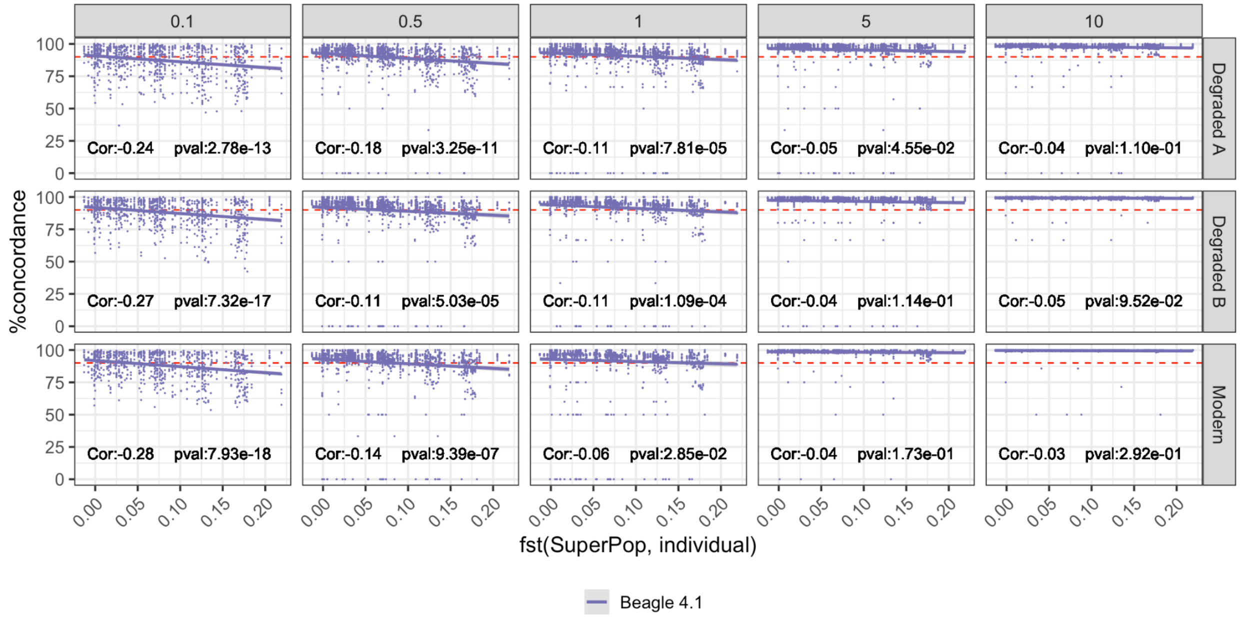


**Figure S19. Correlation between genotype concordance and genetic similarity between an individual and the population reference panel used for the human origins SNP panel.** Data for all tested population reference panels was included. Correlation and p-values were calculated using Pearson's correlation test. Higher $F_{ST}$ values indicate less genetic similarity.

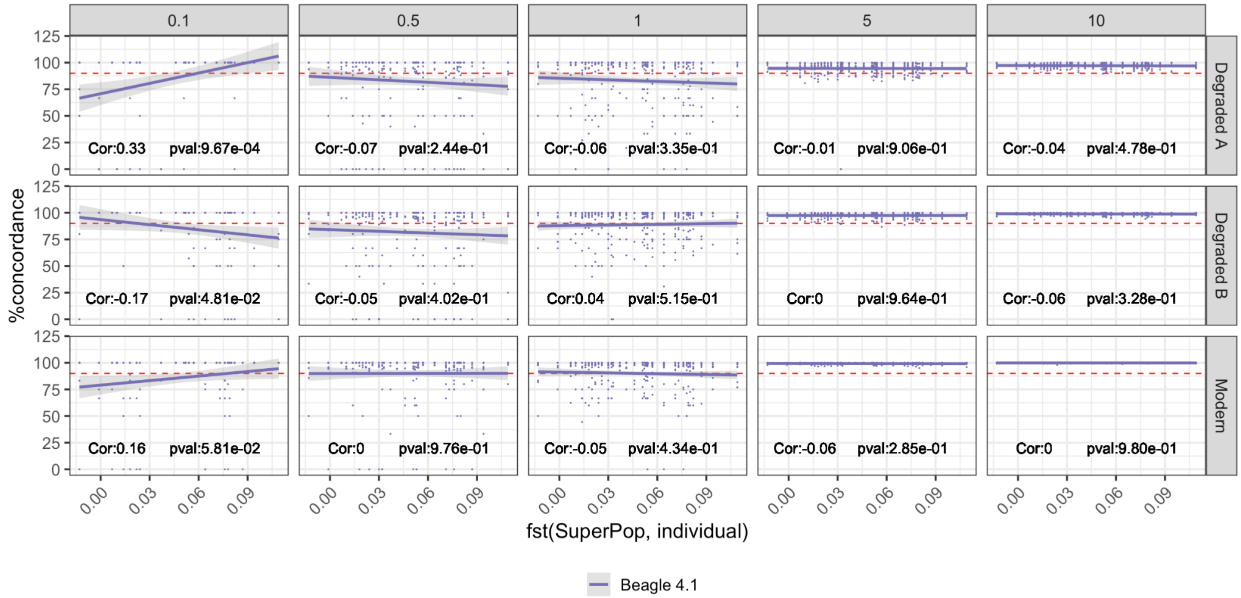


**Figure S20. Correlation between genotype concordance and genetic similarity between an individual and the population reference panel used for the 25K SNP panel.** Data for only the population reference panel of 450 individuals across all five super populations in the 1000 genomes dataset was used. Correlation and p-values calculated using Pearson's correlation test. Higher $F_{ST}$ values indicate less genetic similarity.

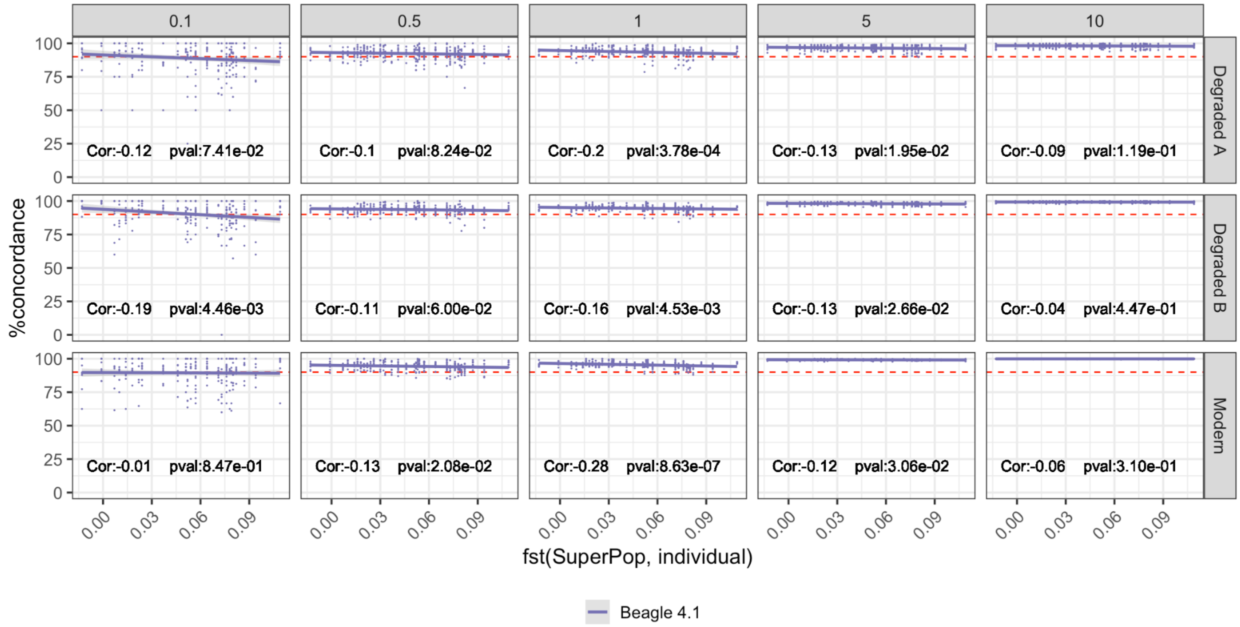


**Figure S21. Correlation between genotype concordance and genetic similarity between an individual and the population reference panel used for the 95K SNP panel.** Data for only the population reference panel of 450 individuals across all five super populations in the 1000 genomes dataset was used. Correlation and p-values calculated using Pearson's correlation test. Higher $F_{ST}$ values indicate less genetic similarity.

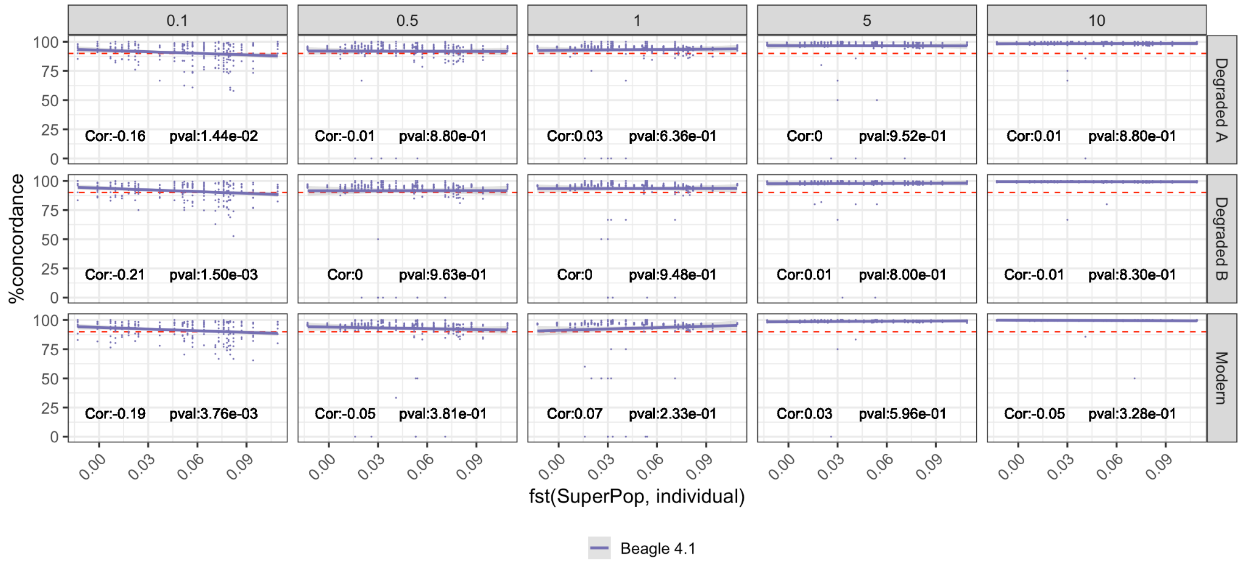


**Figure S22. Correlation between genotype concordance and genetic similarity between an individual and the population reference panel used for the human origins SNP panel.** Data for only the population reference panel of 450 individuals across all five super populations in the 1000 genomes dataset was used. Correlation and p-values calculated using Pearson's correlation test.

### 5. Imputation

We investigated the performance of widely used imputation methods including SAMtools workflow with Beagle 5.4(Browning et al., 2018) and GLIMPSE2(Rubinacci et al., 2021, 2023) according to the recommendations for each respective method (Methods 2.6). We used the outputs of the ATLAS genotyping + Beagle 4.1 workflow. We used four different population reference panels, each consisting of 450 individuals. $F_{ST}$ was calculated to describe the genetic affinity between each respective individual and population reference panel (Methods 2.2).

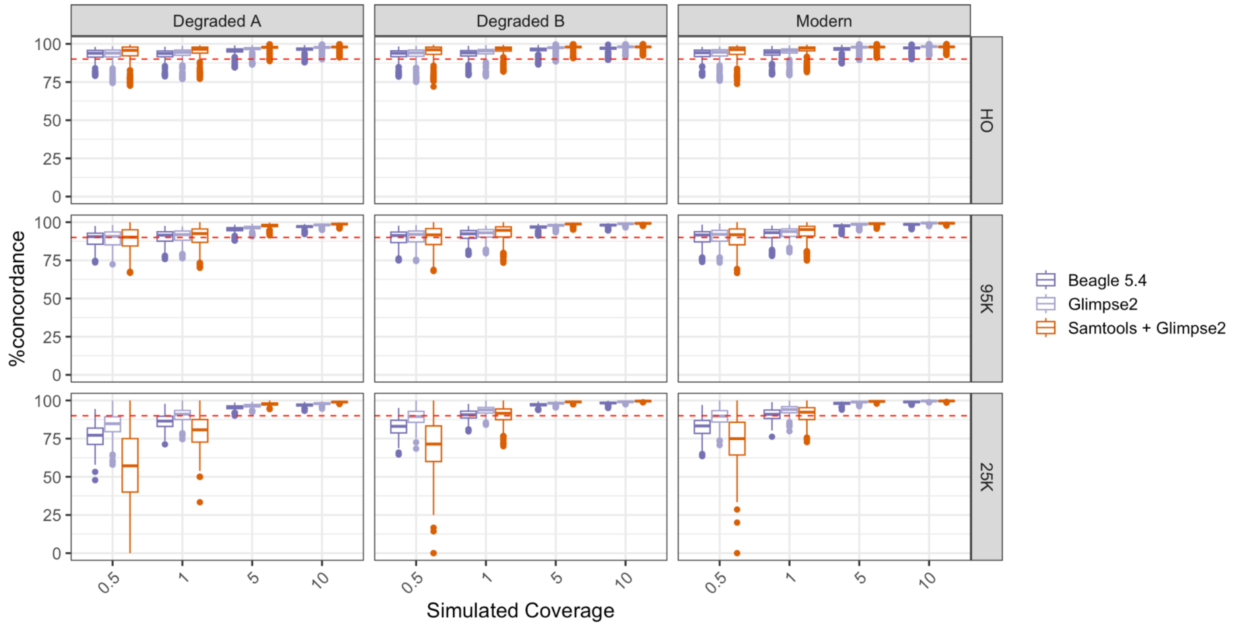


**Figure S23. Concordance of genotypes for different workflows of genotyping and imputation methods with all tested population reference panels.** Genotype likelihood threshold of 99% was used for refinement and imputation. The Tukey box and whisker plots show the lower and upper quartiles boxed with the whiskers representing 1.5 times the interquartile range. Dots denote any outliers. Beagle 5.4: Atlas genotyping + Beagle 4.1 refinement + Beagle 5.4 imputation; GLIMPSE2: Atlas genotyping + Beagle 4.1 refinement +GLIMPSE2

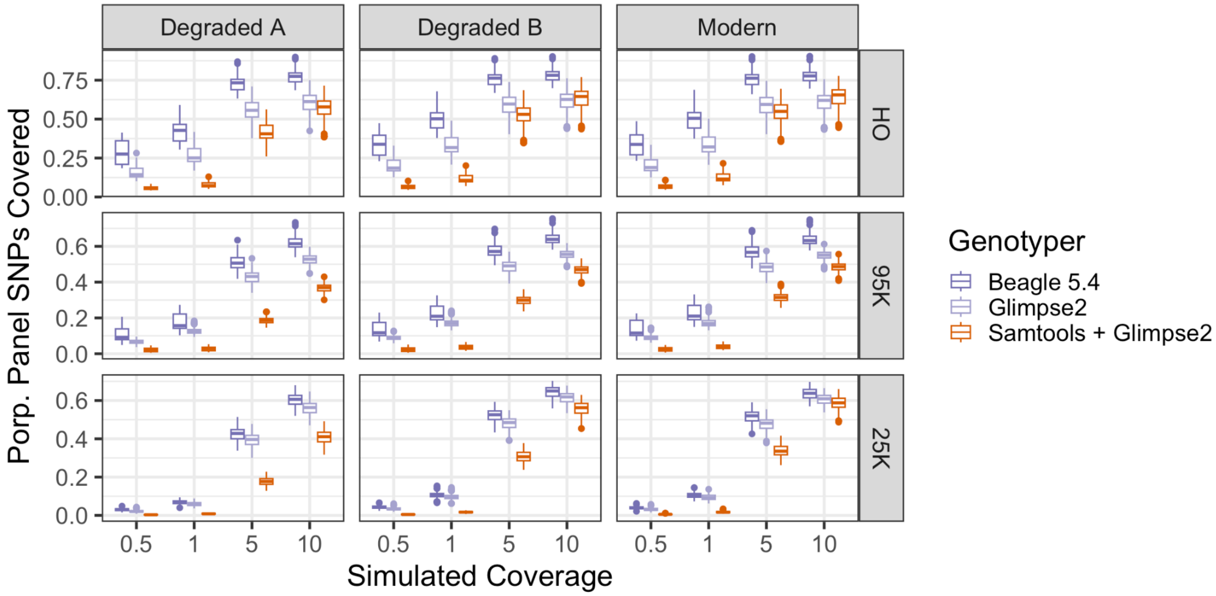


**Figure S14. Proportion of Genotyped SNPs after Imputation for different SNP panels**. Genotype likelihood thresholds of 99% were used for refinement and imputation. Data from all for tested population reference panels is included in the figure. The Tukey box and whisker plots show the lower and upper quartiles boxed with the whiskers representing 1.5 times the interquartile range. Dots denote any outliers. Beagle 5.4: Atlas genotyping + Beagle 4.1 refinement + Beagle 5.4 imputation; GLIMPSE2: Atlas genotyping + Beagle 4.1 refinement +GLIMPSE2

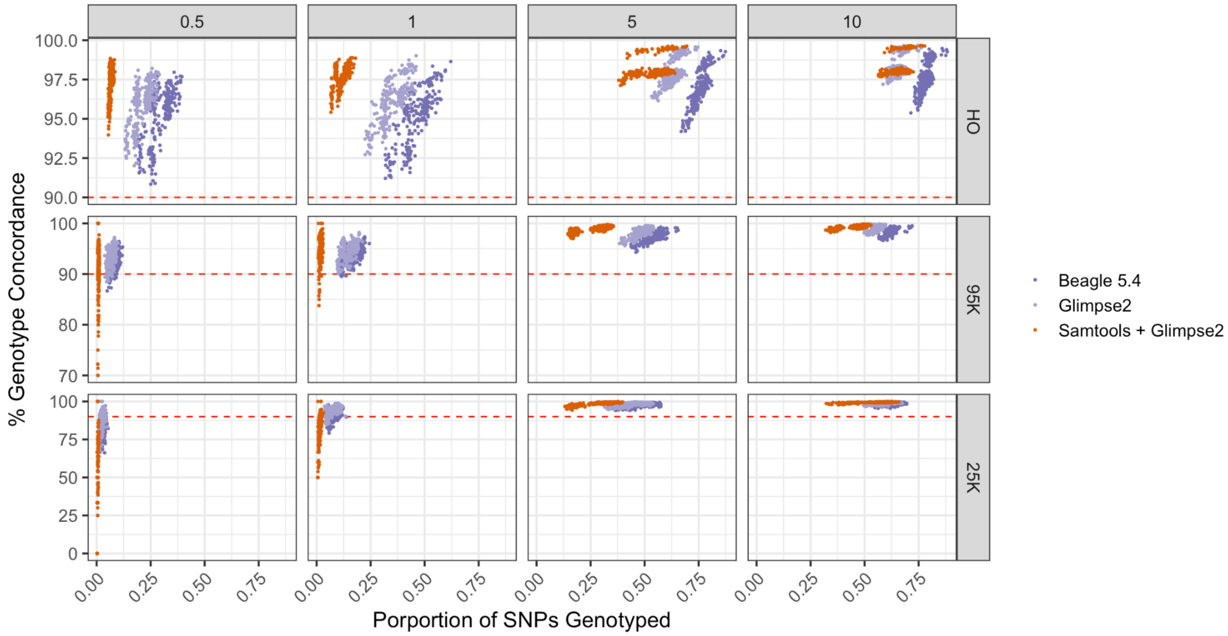


**Figure S25. Correlation between genotype concordances and proportion of genotyped SNPs in a SNP panel after imputation.** Genotype likelihood thresholds of 99% were used for refinement and imputation. Data shown was imputed using a population reference panel representative of the 1000 genomes population reference panel. Red dotted line indicated 90%. Beagle 5.4: Atlas genotyping + Beagle 4.1 refinement + Beagle 5.4 imputation; GLIMPSE2: Atlas genotyping + Beagle 4.1 refinement +GLIMPSE2. Note that the y-axis is different for each panel.

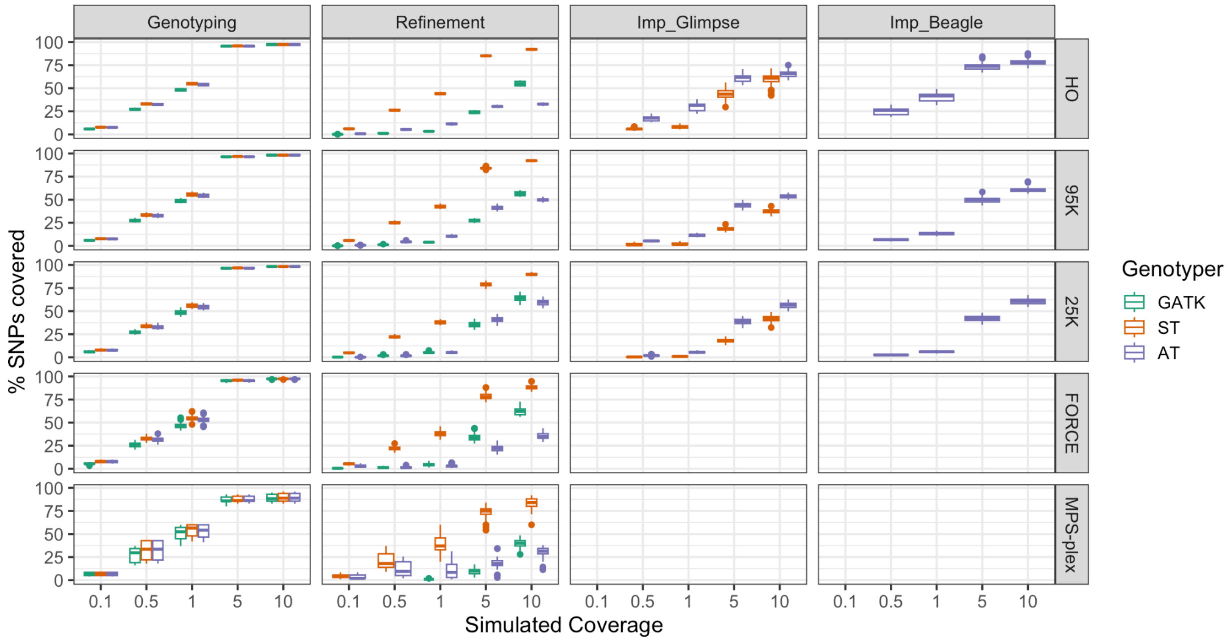


**Figure S26. Percent of SNPs genotyped in each respective panel through the different analysis steps.** Box plots are colored based on which method was used for the initial genotyping. Genotype likelihood thresholds of 99% were used for refinement and imputation. Data shown was imputed using a population reference panel representative of the 1000 genomes population reference panel. Imp_Glimpse= Imputation performed with GLIMPSE2, Imp_Beagle=Imputation performed with Beagle 5.4.

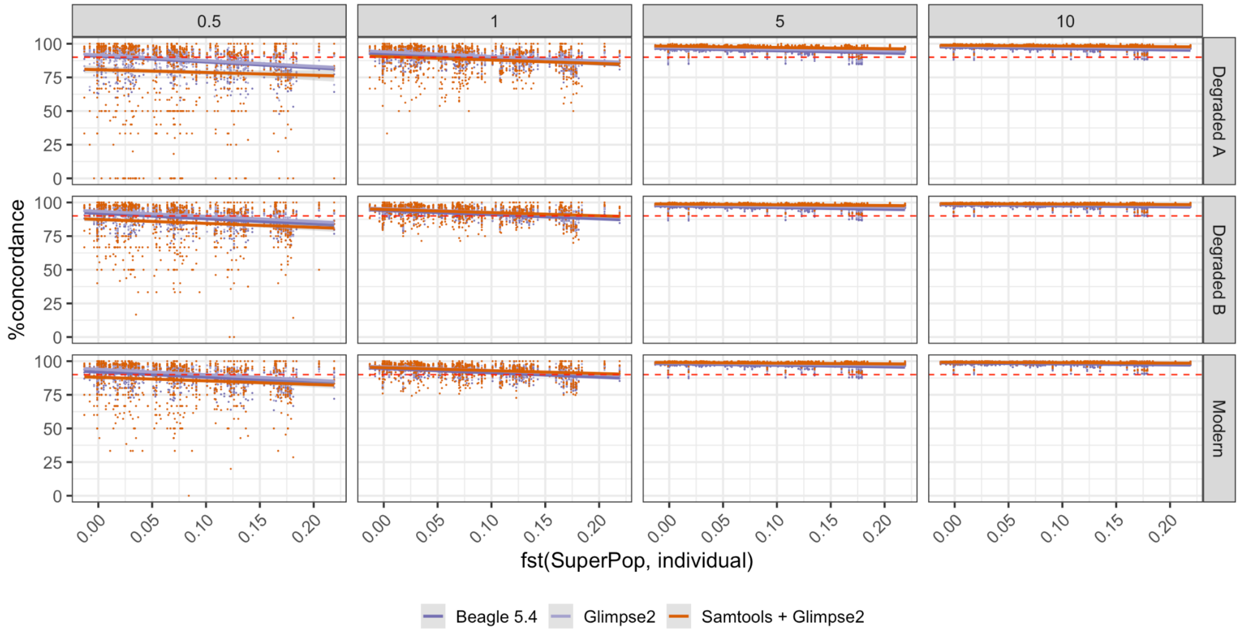

**Figure 27: Correlation between genotype concordance and genetic similarity between an individual and the population reference panel across the HO, 95K, and 25K SNP panels.** Data for all four tested population reference panels (African, East Asian, American, and 1000 genomes) was used. Genotype likelihood thresholds of 99% were used for refinement and imputation. Beagle 5.4: Atlas genotyping + Beagle 4.1 refinement + Beagle 5.4 imputation; GLIMPSE2: Atlas genotyping + Beagle 4.1 refinement +GLIMPSE2.

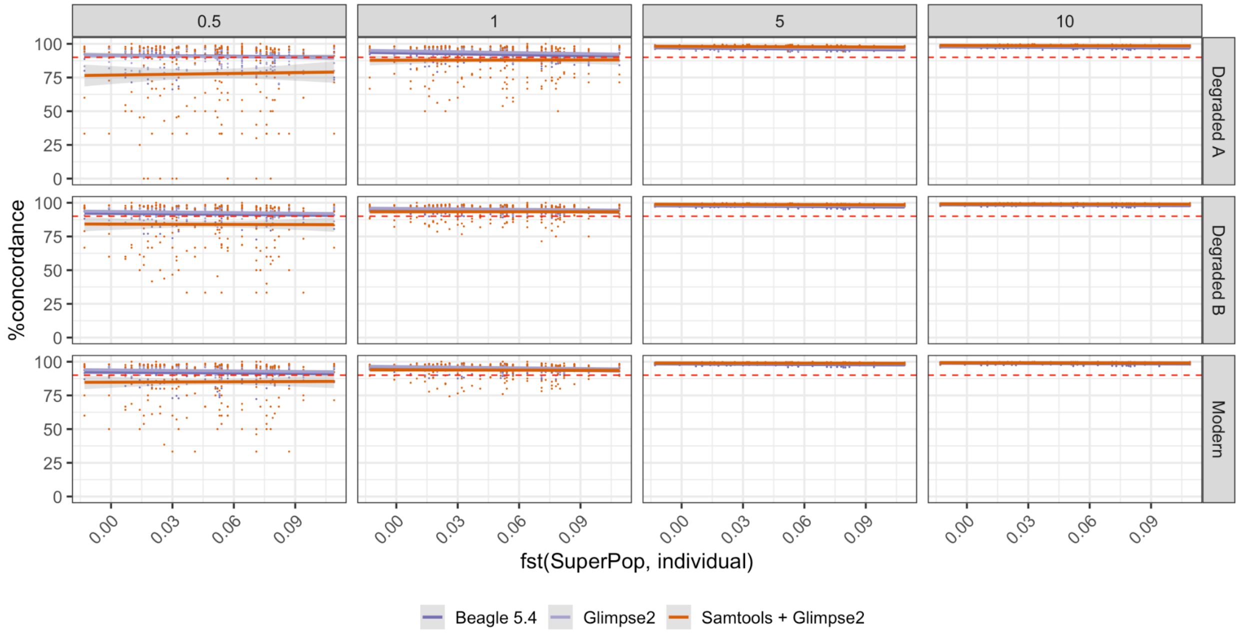


**Figure S28: Correlation between genotype concordance and genetic similarity between an individual and the population reference panel across the HO, 95K, and 25K SNP panels.** Data for only the population reference panel of 450 individuals across all five super populations in the 1000 genomes dataset was used. Genotype likelihood thresholds of 99% were used for refinement and imputation. Beagle 5.4: Atlas genotyping + Beagle 4.1 refinement + Beagle 5.4 imputation; GLIMPSE2: Atlas genotyping + Beagle 4.1 refinement +GLIMPSE2.

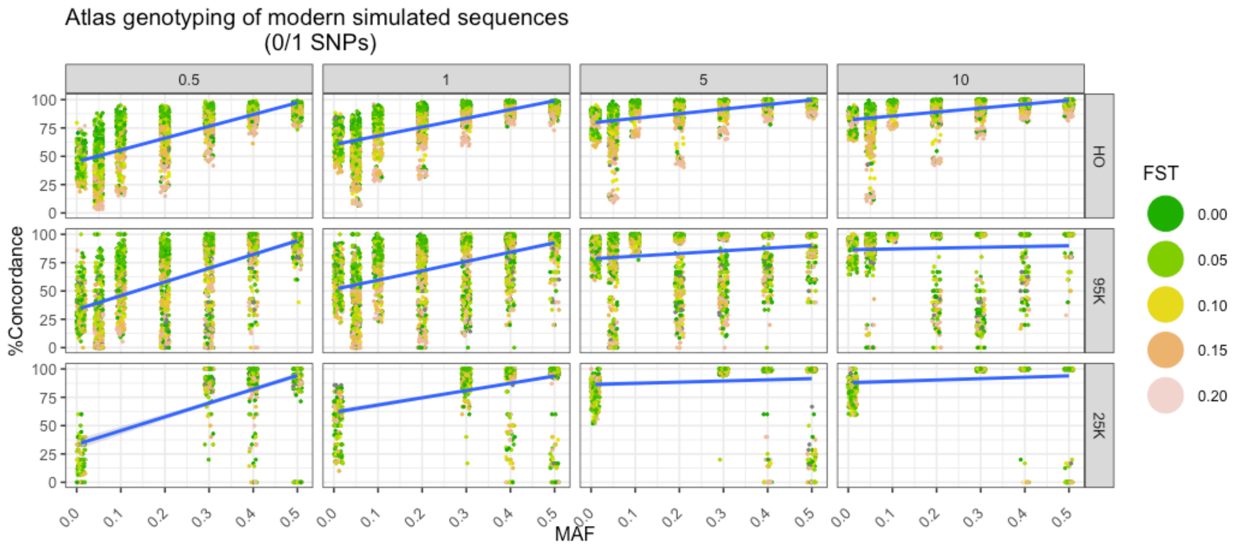


**Figure S29: Correlation between genotype concordance of heterozygous sites and minor allele frequencies (MAF).** Genotype concordance is based on the outputs from genotyping using ATLAS. Data for all four tested population reference panels (African, East Asian, American, and 1000G mixed) was used. Vertical facets represent different simulated coverages. The blue lines are based on a linear model and the gray shaded area represents 95% confidence intervals. MAFs are as follows: 0.01=0-0.01, 0.05=0.011-0.05; 0.1=0.051-0.1; 0.2=0.11-0.2; 0.3=0.21-0.3; 0.4=0.31-0.4; 0.5=0.41-0.5

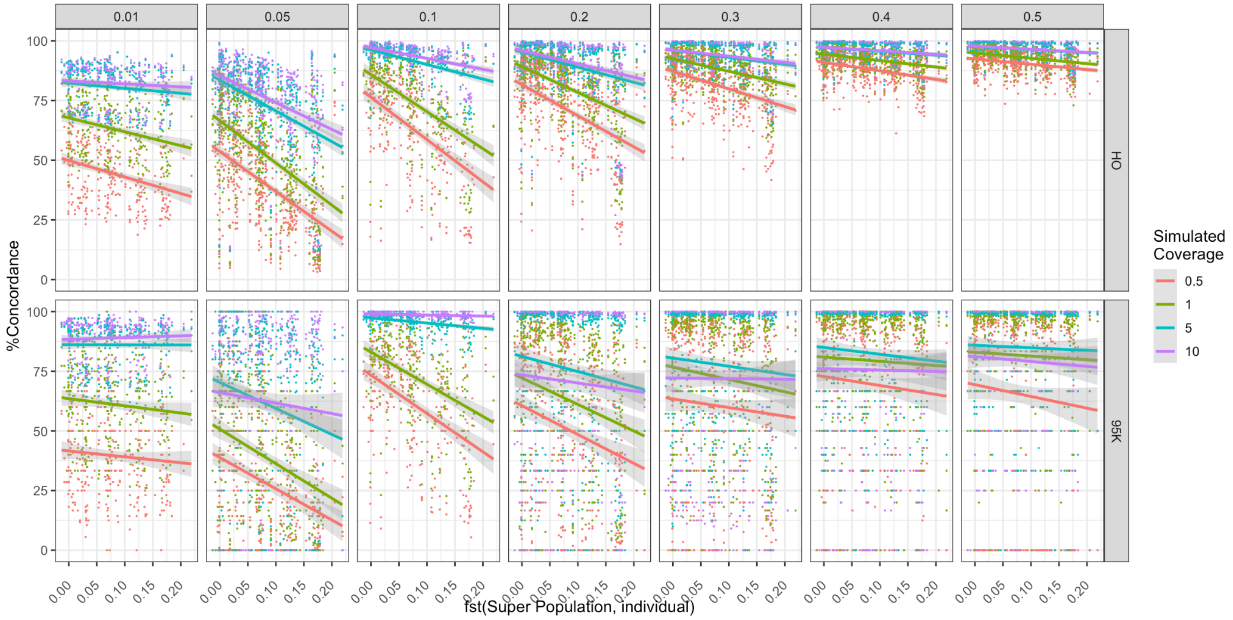


**Figure S30: Correlation between genotype concordance of heterozygous sites and genetic similarity of an individual to the population reference panel across minor allele**

**frequencies (MAF).** The vertical facets are MAFs. Genotype concordance is based on the outputs from genotyping using ATLAS. Data for all four tested population reference panels (African, East Asian, American, and 1000 genomes) was used. The lines are based on a linear model and the gray shaded area represents 95% confidence intervals. MAFs are as follows: 0.01=0-0.01, 0.05=0.011-0.05; 0.1=0.051-0.1; 0.2=0.11-0.2; 0.3=0.21-0.3; 0.4=0.31-0.4; 0.5=0.41-0.5

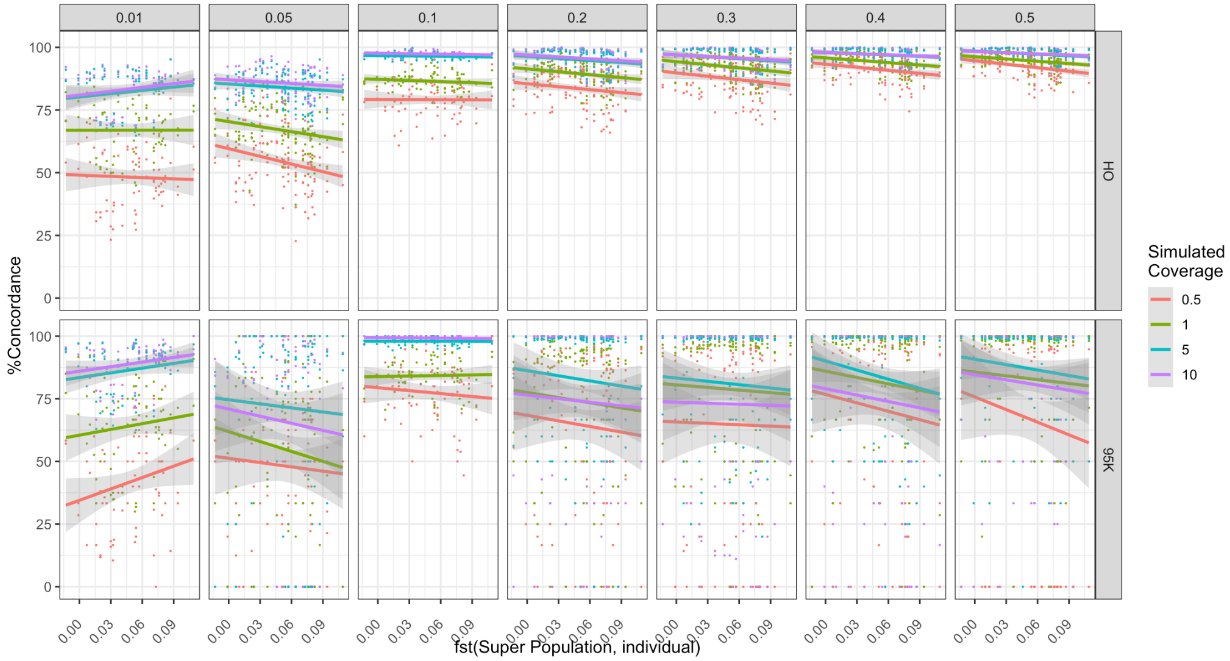


**Figure S31: Correlation between genotype concordance of heterozygous sites and genetic similarity to a 1000 genomes population reference panel across minor allele frequencies (MAF).** The vertical facets are MAFs. Genotype concordance is based on the outputs from genotyping using ATLAS. Data for only the population reference panel of 450 individuals across all five super populations in the 1000 genomes dataset was used. The lines are based on a linear model and the gray shaded area represents 95% confidence intervals. MAFs are as follows: 0.01=0-0.01, 0.05=0.011-0.05; 0.1=0.051-0.1; 0.2=0.11-0.2; 0.3=0.21-0.3; 0.4=0.31-0.4; 0.5=0.41-0.5
